# Gene module reconstruction elucidates cellular differentiation processes and the regulatory logic of specialized secretion

**DOI:** 10.1101/2023.12.29.573643

**Authors:** Yiqun Wang, Jialin Liu, Lucia Y. Du, Jannik L. Wyss, Jeffrey A. Farrell, Alexander F. Schier

## Abstract

During differentiation, cells become structurally and functionally specialized, but comprehensive views of the underlying remodeling processes are elusive. Here, we leverage scRNA-seq developmental trajectories to reconstruct differentiation using two secretory tissues as a model system – the zebrafish notochord and hatching gland. First, we present an approach to integrate expression and functional similarities for gene module identification, revealing dozens of gene modules representing known and newly associated differentiation processes and their temporal ordering. Second, we focused on the unfolded protein response (UPR) transducer module to study how general versus cell-type specific secretory functions are regulated. By profiling loss- and gain-of-function embryos, we found that the UPR transcription factors *creb3l1*, *creb3l2*, and *xbp1* are master regulators of a general secretion program. *creb3l1/creb3l2* additionally activate an extracellular matrix secretion program, while *xbp1* partners with *bhlha15* to activate a gland-specific secretion program. Our study offers a multi-source integrated approach for functional gene module identification and illustrates how transcription factors confer general and specialized cellular functions.

## Introduction

In metazoan development, different cell types emerge through the process of differentiation. During differentiation, cellular features are remodeled to confer specific structural and functional properties. Some examples of remodeling include the expansion of particular organelles, alteration of the cell cycle, repurposing of the cytoskeleton, change of adhesive properties, and the creation of new structures. While many genes that underlie these remodeling processes have been identified, such as for cilia or sarcomere assembly, our knowledge of the genes and their associated functions that drive differentiation remains fragmentary^1–6^. Furthermore, while key transcription regulators have been identified, their target genes are often undefined and frequently masked by regulatory redundancy.

The advent of large-scale single-cell RNA sequencing (scRNA-seq) creates the opportunity to analyze biological processes at comprehensive scales. In particular, developmental trajectories reconstructed from scRNA-seq data enable the inference of gene expression cascades that accompany differentiation, which includes the transcription factors (TFs) that may regulate differentiation effector genes^7–12^. While commonly used to define cell type markers, the potential of these trajectories in uncovering cellular remodeling processes during differentiation remains less explored^7^.

Here, we leverage the reconstructed scRNA-seq transcriptional trajectories to build a full compendium of differentiation genes and their associated remodeling processes for given cell types. As model systems, we use two secretory tissues – the zebrafish notochord and hatching gland. The notochord secretes a collagen-rich extracellular matrix (ECM) to enclose vacuolated cells to form a stiff rod, while the hatching gland secretes enzymes that digest the chorion, the protective egg envelope^13,14^. Beyond its characteristic as a hallmark of chordates, the notochord is also involved in patterning neighboring tissues^14^. The notochord and hatching gland cells provide ideal model systems to study differentiation: within the first 20 hours of embryogenesis, they derive from the Mangold-Spemann organizer, form the axial mesoderm, and differentiate into highly specialized cells with prototypical secretory structures and functions (**Figure 1A**)^8^.

**Figure 1.**
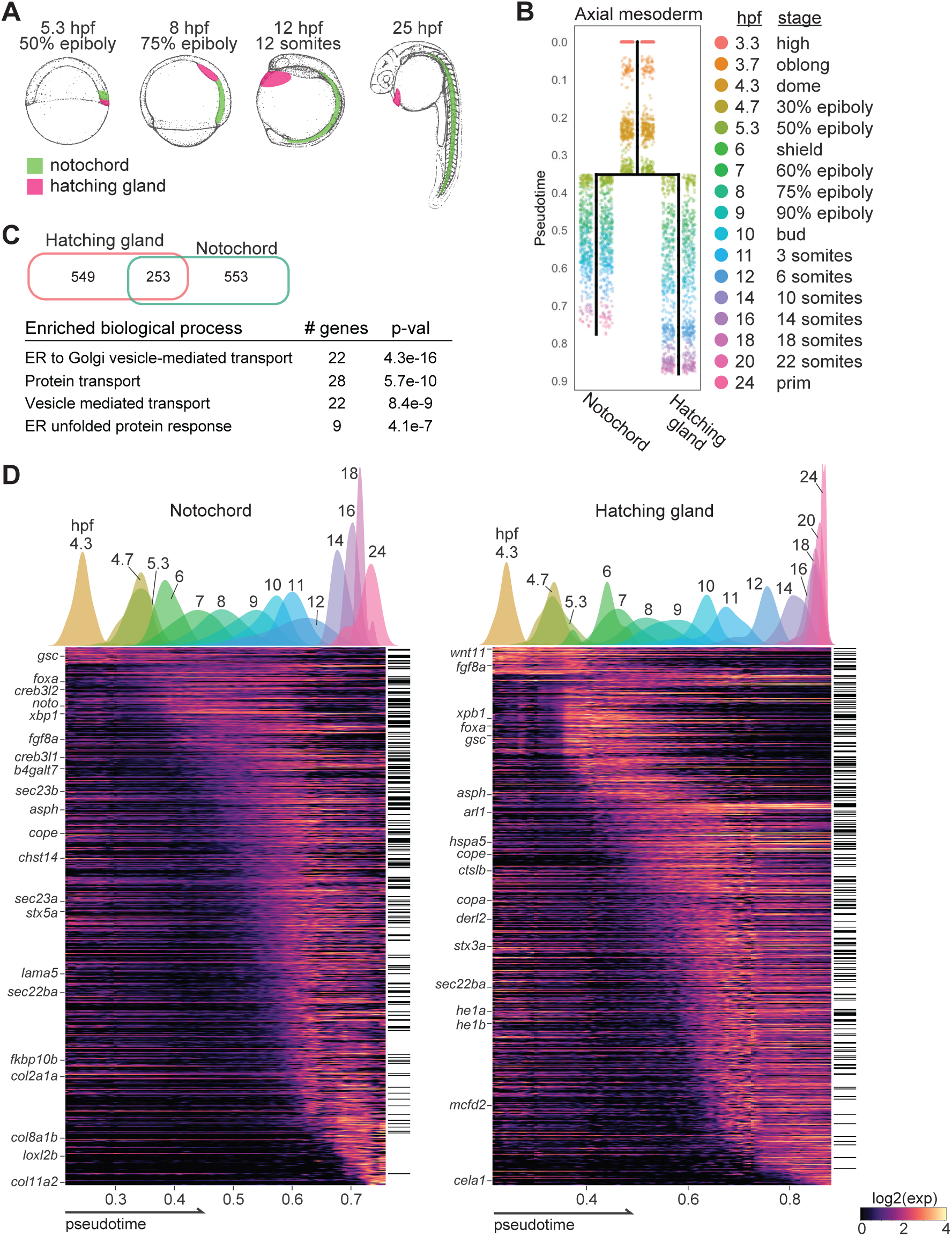
Developmental trajectories reveal hundreds of genes expressed during axial mesoderm differentiation. **A)** Overview of notochord and hatching gland development in the zebrafish embryo. **B)** Transcriptional trajectories of the notochord and the hatching gland reconstructed from scRNA-seq data spanning 3.3 to 24hpf. Points represent single cell transcriptomes colored by stages. Pseudotime measures developmental progression. **C)** Venn diagram of enriched genes shows a substantial overlap between the two cell types (upper). Gene Ontology (GO) terms related to secretion are most statistically enriched in the shared genes. **D)** Expression heatmaps of genes enriched in the notochord (left) and the hatching gland (right). Black bars on the right mark the genes enriched in both cell types. Plots on top show the distribution of developmental stages along pseudotime. Selected genes in pathways discussed later in the main text are labeled on the left.

Past studies of zebrafish axial mesoderm development have identified genes involved in remodeling processes ranging from cell migration, adhesion, and polarization, to ECM secretion, vacuolation and secretory granule formation^13–19^. These landmark studies have revealed essential aspects of notochord and hatching gland formation, but the described genes account for only a fraction of those upregulated during axial mesoderm differentiation, leaving a full picture of the differentiation processes elusive. In addition, the gene regulatory logic dictating the differentiation of distinct secretory cell types is not well understood. For example, the notochord and hatching gland secrete very different cargos: large ECM proteins in the notochord and small proteases in the hatching gland^14,20^. It is poorly understood how such secretory cells meet both shared and cell-type specific secretory challenges. In particular, transcription factors (TFs) in the Unfolded Protein Response (UPR) have been implicated in axial mesoderm differentiation, yet their redundancy, specificity, and target genes are poorly defined^17,18,21,22^. Specifically, are these related factors interchangeable between the two cell types, or are they specialized in activating distinct notochord- or hatching gland-specific programs?

To investigate these questions, we developed an approach that integrates single-cell developmental trajectories with protein interaction and functional annotation databases to identify modules of genes with shared temporal expression and functions. We used these modules to represent remodeling processes and describe their temporal ordering during differentiation. We further investigate the regulation of general and specialized secretory functions by genetically disrupting or mis-expressing candidate TFs and performing scRNA-seq phenotyping. Our results provide a comprehensive characterization of cellular processes during notochord and hatching gland differentiation, with over 800 genes categorized into 70 functional gene modules in each tissue, and reveal the regulatory logic by which different UPR pathway TFs activate shared and specialized secretion machineries. More broadly, our study establishes a generalizable approach to dissect differentiation at genome-wide scale.

## Results

### Transcriptional trajectories identify 1,355 genes with enriched expression in the differentiating notochord and hatching gland

To uncover the genes involved in notochord and hatching gland differentiation, we first sought to systematically identify the genes expressed during their development. We computationally isolated the axial mesoderm cells from our published scRNA-seq datasets based on known marker gene expression and used the URD package to reconstruct their transcriptional trajectories from 3 hours post-fertilization (hpf) to 24hpf (**Figure 1B**)^8,23^. One cell type was identified at 3hpf to serve as the progenitor cell type, while 2 cell types (1 notochord and 1 hatching gland) were identified at 24hpf, which served as the terminal cell types for the URD algorithm. The reconstructed trajectories represent the transcriptional progression from the progenitor state towards the terminal, differentiated states.

Cells were assigned pseudotime based on their transcriptional distance from the progenitor state, which provides a continuous metric of a cell’s developmental progression. We divided each transcriptional trajectory into 15 approximately equal pseudotime windows and defined enriched genes as those with stronger expression in the notochord or hatching gland compared to the rest of the embryo from the same pseudotime windows (Methods). This approach identified over 800 enriched genes per cell type, including 253 overlapping genes (**Figure 1C-D**). These shared genes are enriched for secretion-related functions (e.g. vesicle-mediated transport), consistent with the secretory functions of notochord and hatching gland (**Figure 1C**). Expression dynamics revealed that enriched genes are activated in temporal waves, reminiscent of the structural and functional changes during differentiation (**Figure 1D**).

### Integrating transcriptional trajectories and functional annotation databases defines gene modules

To categorize the enriched genes and identify cellular processes underlying notochord and hatching gland development, we first used gene ontology (GO) analysis. This approach identified 220 and 180 biological processes overrepresented (p<0.05) among the enriched genes in the notochord and hatching gland, respectively (**table S1-2**). The most significant GO terms described shared processes within these cell types, such as secretory functions (“vesicle-mediated transport” and “protein folding”), as well as cell-type-specific roles (“collagen fibril organization” in the notochord, and “hatching” in the hatching gland). Despite recovering important biological processes, several factors made GO analysis results difficult to interpret. For instance, highly redundant terms (e.g. “SRP-dependent cotranslational protein targeting to membrane” and “protein targeting to ER”) and spurious terms (e.g. “endoderm development” and “somite development”) were frequently recovered. Moreover, the over-represented biological processes only included about half of the enriched genes in each cell type. Among these genes, ∼35% were associated with at least 3 processes, making their specific functions in axial mesoderm ambiguous.

To group genes into modules representing cellular processes, we explored alternative strategies to leverage multiple sources of data, including expression dynamics and additional functional annotations. To integrate information from scRNA-seq data and functional annotation databases, we calculated genes’ similarities in both expression dynamics and annotated functions. Jensen-Shannon divergence between genes’ expression dynamics was used to measure expression similarities, while functional similarities were calculated from several publicly available functional annotation databases, including STRING, Reactome, GO, and Interpro^24–27^. The gene-gene interaction scores from STRING were directly used as similarity scores. The other databases provided functional annotations for individual genes, and we calculated the semantic similarities between genes’ annotated terms to represent functional similarities^28^. Functional similarities from different sources were then combined (Methods).

We next clustered the genes into modules based on similarity scores, assigning each gene to a unique module. This approach may ignore secondary functions for multi-functional genes but offers two key advantages that enhance the interpretability of the results: first, it avoids ambiguity about the cellular process associated with each gene; and second, it prevents the redundant identification of similar processes containing highly overlapping genes. A range of metrics and parameters were tested, including using expression or functional similarity alone, combining the similarities with different methods, as well as using different clustering algorithms and parameters (Methods). We selected the methods that assigned most genes (>85%) into non-singlet clusters while producing reasonable cluster numbers (<150 clusters) and sizes (<150 genes for the largest cluster) for each cell type. We then manually classified 300 randomly selected notochord-enriched genes into modules based on their expression dynamics and literature, and further calibrated the clustering methods based on their agreement with our manual curation (**Figure S1A-B**, Methods). Based on these criteria, we found the best approach to gene module identification was to cluster genes using the Leiden algorithm based on the product of their expression similarity and functional similarity (**Figure 2A**).

**Figure 2.**
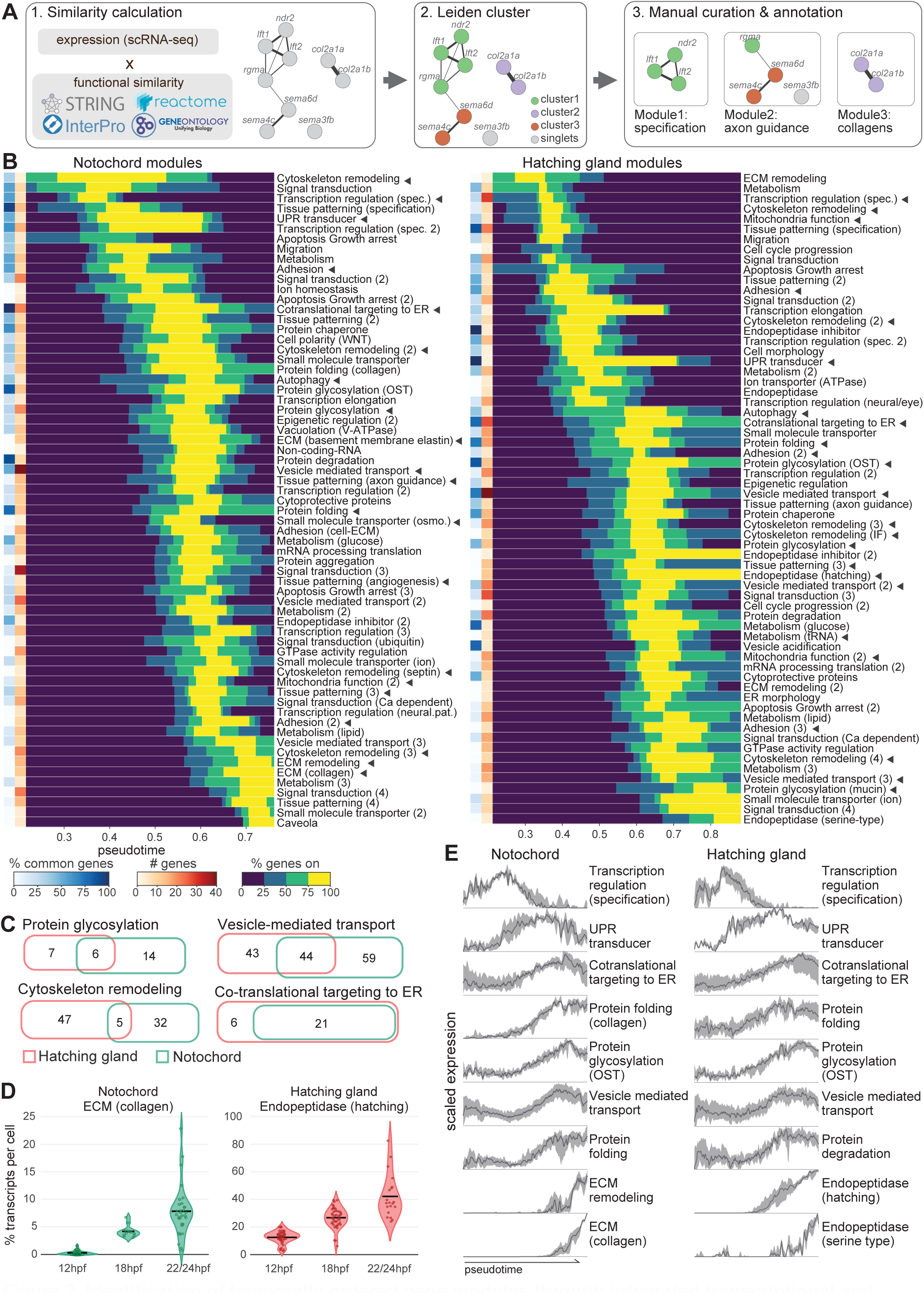
Identification of temporally ordered gene modules through integrated transcriptional and functional similarities. **A)** Schematic of gene module identification workflow. **B)** Cascades of identified gene modules. Rows correspond to modules. Blue side bars indicate the percentage of module member genes commonly enriched in both cell types. Red side bars indicate the number of module member genes. Heatmap is colored by the percentage of genes in each module that is “on” at each pseudotime point. Development progresses from lower to higher pseudotime (horizontal axis). Arrowheads point to the modules mentioned in the text. Modules with similar functions but which are temporally separated share the same names with different numbers appended to the end. Abbreviations: “spec.” - specification; “osmo.” - osmoregulation; “neural.pat.” - neural patterning. **C)** Member gene overlap between similar modules in the two cell types. **D)** Percentage of the secretory cargo coding transcripts per cell at late developmental stages. Percentages are calculated as the sum of transcripts from genes in the “ECM (collagen)” or the “Endopeptidase (hatching)” module divided by the total transcripts per cell. Black horizontal bars mark mean values. **E)** Expression dynamics of selected gene modules. Dark solid line represents median expression among genes in the module at each pseudotime point, while lower and upper bounds of the ribbon indicate the 25 and 75 percentiles.

Notably, clustering with integrated expression and functional similarities outperformed either expression or functional similarities alone (**Figure S1B**). Clustering based solely on expression similarities performed poorly because simultaneous processes were often combined. For instance, a co-expression cluster contained genes that encode developmental TFs (e.g. *foxa1*, *irx3a*)^29,30^, ECM components (e.g. *fn1a*, *lamc1*)^31^, and protein folding enzymes (e.g. *p4ha1a*, *plod2*)^32^. Conversely, using functional similarities alone combined genes that contribute to temporally separated processes. For instance, many secreted factors were combined into one module, despite participating in temporally distinct patterning events, such as patterning the early axial mesoderm (e.g. *admp*, *chd*)^33^, or the tissues that surround the notochord (e.g. *shha*, *slit2*)^34,35^. Therefore, using a combination of transcription trajectories and functional annotation databases best determines functionally and temporally distinct processes. We call this approach MIMIR (after the wise Norse god, or for <u>M</u>odule Identification via <u>M</u>ulti-source Integration for <u>R</u>econstructing Differentiation). Scripts for running MIMIR are available on GitHub (https://github.com/YiqunW/MIMIR).

To produce a final set of modules, we applied MIMIR using the optimized parameters. MIMIR initially generated 86 notochord and 74 hatching gland gene modules that contained at least two genes, covering ∼90% of the enriched genes. These initial modules were then refined, which involved splitting or merging modules to ensure coherent gene functions and expression dynamics within them (Methods). By manually checking annotations, expression dynamics, and literatures, we additionally assigned 88 previously excluded genes to modules, resulting in the final assignment of 97% notochord-enriched genes into 71 modules and 95% hatching gland-enriched genes into 70 modules (**Figure 2A-B**, **table S3-4**). Comparison between automated and curated modules yielded an adjusted mutual information (AMI) score of 0.66 and 0.67 for notochord and hatching gland, respectively, where 0 represents random assignment and 1 represents a perfect match (**Figure S1C**). Modules were then manually annotated according to their member gene functions, based on functional annotations and literature (Methods). Compared to conventional GO enrichment analysis, MIMIR uncovered more non-redundant processes and assigned unambiguous functions to >500 more genes in each cell type. In conclusion, our approach combines transcriptional trajectories and functional annotation databases to comprehensively assign genes enriched in differentiation trajectories into temporally resolved functional gene modules.

### Gene module analysis reveals known and novel cellular processes

Our goal in constructing gene modules was to identify cellular processes underlying notochord and hatching gland differentiation. Inspection of the gene modules confirmed that many modules reflect known cell type-specific functions. For instance, the notochord is known for producing a collagen-rich ECM sheath; accordingly, we found two ECM component modules specifically in the notochord, one contains basement membrane components (e.g. *lama5*) and the other collagens (e.g. *col2a1a*)^14^.

Similarly, the hatching gland is known for producing hatching enzymes, and an endopeptidase module containing conserved hatching enzymes (e.g. *he1a*) was specifically found in the hatching gland^20^. In contrast, some modules represent shared processes known to occur in both cell types. For example, both cell types expressed mesendoderm specification modules containing well-known TFs (e.g. *foxa3*, *gsc*, *sebox*), as well as modules for signal transduction, cytoskeleton remodeling, and vesicle-mediated transport, among others^36,37^. While representing shared processes, these modules often have different, cell type-specific genes. For instance, the cytoskeleton remodeling module in the hatching gland includes intermediate filament components (e.g. *krt18*, *krt8*) that are absent from the notochord; the protein glycosylation module in the notochord contains enzymes with specialized roles in modifying ECM proteoglycans not found in the hatching gland (e.g. *chst14*, *chst2a*) (**Figure 2C**)^32^. In contrast, only a few shared modules between the two cell types contain highly overlapping genes, such as co-translational targeting to ER (**Figure 2B-C**). Thus, effector genes driving similar biological processes were often fine-tuned in different cell types to address cell-type-specific needs.

Both cell-autonomous and non-autonomous functions were recovered as modules. Non-autonomous modules were named “Tissue patterning”, as they contain signaling genes that can instruct the fate or behavior of nearby cells. Some examples are the notochord tissue patterning modules that include angiogenesis signals (e.g. *angptl6*, *vegfaa*)^38^, axon guidance signals (e.g. *sema4c*, *slit2*)^39^, and floor plate induction signals (e.g. *shha*)^40^. Thus, modules in a cell type not only identify processes essential for differentiation of that cell type but also reveal effects on other cell types.

While many genes in these modules were well-known, some were not well described. For example, the collagen module in the notochord includes nine well-annotated collagen genes (e.g. *col2a1a*, *col9a2*, *col11a2*), as well as two previously uncharacterized genes (*cabz01044297.1*/*zmp:0000000760*, *zgc:113232*) that share predicted collagen triple helix structure (based on Interpro) and similar expression dynamics with the other collagens in the module. Similarly, the notochord vacuolation module contains several vacuolar-type H(+)-ATPase (V-ATPase) subunits, some with previously demonstrated roles in notochord vacuolation, such as *atp6v1e1b*, and some with undescribed notochord function, such as *atp6v0e1*^16^. Using CRISPR-Cas9, we verified the requirement for *atp6v0e1* in notochord vacuolar structure and revealed its functional redundancy with *atp6v1e1b* (**Figure S2**). MIMIR therefore helps associate under-characterized genes with biological functions during differentiation.

We also identified novel modules that represent processes previously unassociated with the notochord and hatching gland differentiation. For instance, both cell types expressed a module of genes that function within mitochondria (e.g. *mthfd2*, *uqcc3*) and a module of genes involved in autophagosome formation (e.g. *gabarapl1*). Such modules may satisfy these cells’ demands in high energy consumption, protein quality control, and non-canonical secretory processes^40^.

Some novel modules are specific to either notochord or hatching gland. For example, a tRNA processing module in the hatching gland (e.g. *tsen34*, *vars*) may reflect high demand for tRNA by the massive production of hatching enzymes (**Figure 2D**). Another example is the notochord-specific module of small molecule transporters responsive to osmotic changes (e.g. *lrrc8aa*, *lrrc8da*)^41^, which could potentially modulate cellular and/or vacuolar osmotic pressure during notochord vacuolation.

In summary, our identified modules recapitulated known processes, identified shared and differential gene usage between cell types for shared processes, associated under-characterized genes, and proposed novel processes in notochord and hatching gland differentiation. These results demonstrated the usefulness of MIMIR in identifying differentiation processes and in generating new hypotheses for exploring functions of understudied genes.

### Temporal order of gene modules uncovers differentiation dynamics

To reveal the temporal progression of differentiation processes, we analyzed the sequence of module expression within our transcriptional trajectories. The expression onset and offset times for each enriched gene were inferred by fitting an impulse model to gene expression along pseudotime in each cell type trajectory^8^. We then ordered the modules according to the average onset times of their member genes and plotted the percentage of active genes within each module at each timepoint to visualize their activity duration (**Figure 2B**).

Inspecting module cascades revealed that the two cell types share several major themes in their cellular remodeling dynamics. For instance, in both cell types, development starts with the expression of cell fate specification modules; afterwards, cells express waves of adhesion and cytoskeleton remodeling modules (**Figure 2B**). These observations are consistent with our understanding that the notochord and hatching gland alter their adhesive properties and morphologies while migrating during differentiation^15,42^. After specification, both cell types also activate modules for their secretory machineries, including co-translational targeting to the ER, protein folding, protein glycosylation, and vesicle-mediated transport (**Figure 2E**). Finally, modules of secretory cargos (i.e. collagens in the notochord and endopeptidase in the hatching gland) turn on and remain active at 24hpf (**Figure 2E**). Thus, module dynamics reveals the sequential biological processes underlying cellular specification and differentiation.

The temporal order of module expression suggested that developmental programs anticipate future needs of differentiating tissues. For instance, a protein folding module that specializes in folding collagens in the ER is expressed before the collagen module in the notochord (**Figure 2E**). Similarly, a module of UPR transducers (e.g. *xbp1*, *hspa5*) is expressed in each cell type before cargo protein production (**Figure 2E**). The UPR is a highly conserved ER stress response induced by unfolded or misfolded proteins to expand the secretory machinery of cells^43,44^. Cells that produce high levels of secretory proteins are prone to such ER stress^45^. In late notochord and hatching gland cells, secretory cargo protein transcripts make up on average 8% and 40% of total transcripts per cell, respectively (**Figure 2D**). The early remodeling of the secretory machinery in advance of cargo production indicates that future secretory functions are anticipated in notochord and hatching gland cells.

### The UPR transducer module contains general and cell type-specific transcription factors

Having defined the composition and dynamics of differentiation modules, we next investigated the gene regulatory mechanisms governing shared and cell type-specific cellular processes. For example, the same master TF might be active in different cell types and activate the same shared differentiation program. This scenario is at play in the RFX-mediated activation of the ciliary program in different neurons^2,46^. Alternatively, in a process of phenotypic convergence, different cell type-specific factors might activate shared differentiation programs in different cell types^47,48^. For example, cholinergic properties are controlled by different transcription factors in different neurons^49^. To explore these and other regulatory strategies, we focused on the UPR transducer modules as a model system. The UPR has both widely conserved and specialized roles in cellular remodeling^45,50–53^. We found that the UPR transducer module member genes are specifically enriched in the axial mesoderm trajectories while showing low or no expression in other cell types in the 3-12hpf scRNA-seq data (**Figure 3A**, **Figure S3**). The UPR transducer modules contain genes shared between the two cell types as well as notochord-specific genes (**Figure 3B**). Both cell types express the chaperone *hspa5*/*bip*, which monitors unfolded and mis-folded proteins in the ER, and UPR TFs *xbp1* and *atf6*, which activate downstream genes upon unfolded protein detection. The notochord UPR transducer module contains two additional UPR TFs (*creb3l1* and *creb3l2*), as well as *mbtps1*, a protease required for the post-translational activation of Atf6, Creb3l1, and Creb3l2^54^. The observation that both shared and notochord-specific UPR transducers are expressed during notochord and hatching gland differentiation and raise the question how the UPR TFs are deployed to confer shared and specific secretory functions.

**Figure 3.**
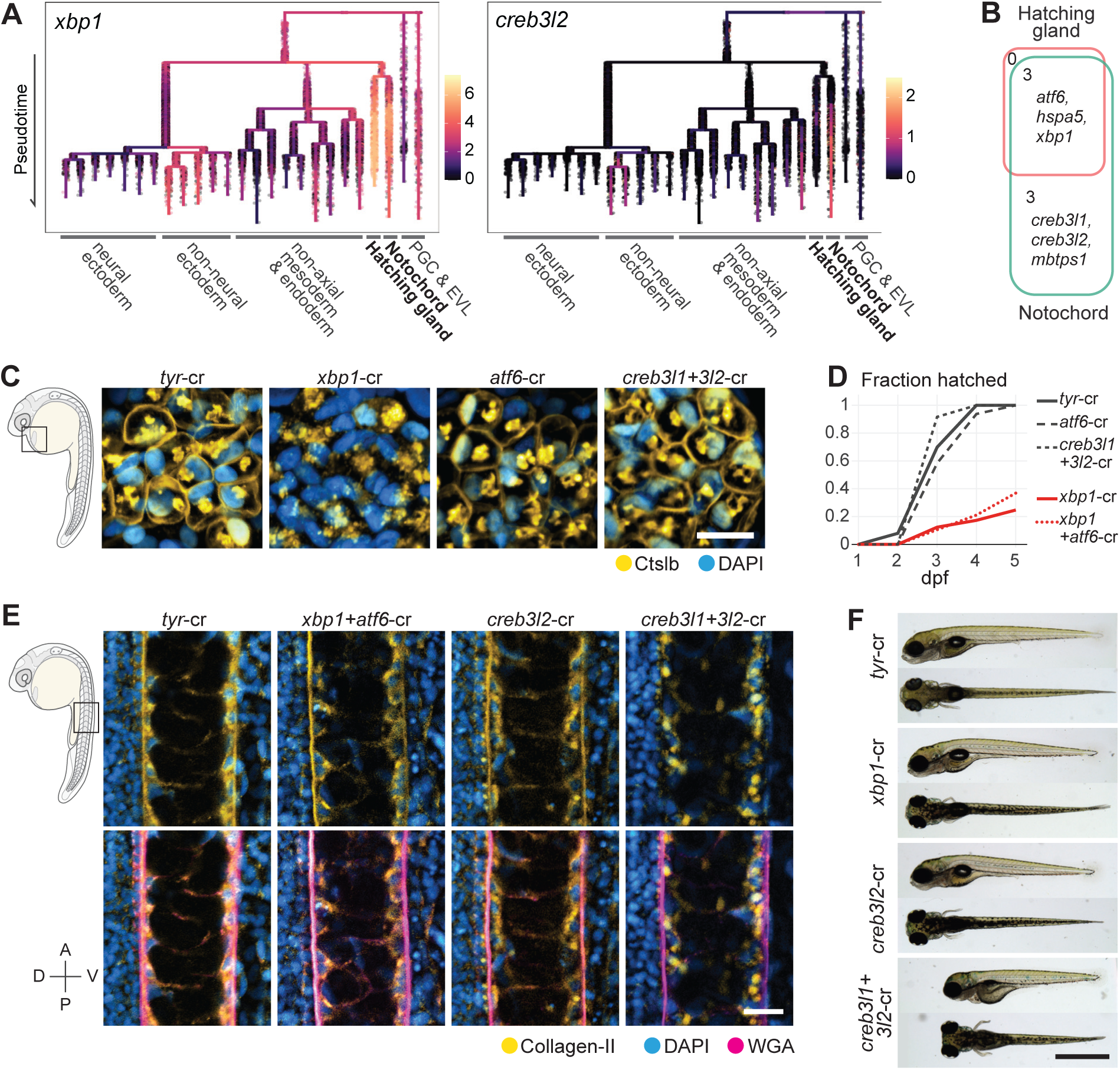
Transcription factors involved in unfolded protein response are required for hatching gland and notochord differentiation. **A)** Expression of a shared (*xbp1*) and a notochord-specific (*creb3l2*) UPR transducer during early zebrafish embryogenesis. Dendrogram represents developmental trajectories of embryonic cell types from 3 to 12hpf (top to bottom of the dendrogram). Dots are single cell transcriptomes colored by log gene expression. See Figure S3 for the expression of *atf6*, *creb3l1*, *mbtps1*, and *hspa5*. **B)** Venn diagram and identities of UPR transducer module member genes in the two cell types. **C)** Immunofluorescence of Ctslb. Images are of hatching gland cells in 28hpf embryos. Loss of *xbp1* resulted in reduced Ctslb on the cell surface. Blue: DAPI nuclei stain; yellow: Ctslb. Scale bar: 20um. D) *xbp1* crispants exhibited reduced hatching. Number of embryos assayed for each condition: tyr-cr=63; *atf6*-cr=48, *creb3l1+3l2*-cr=12; *xbp1*-cr=81; *xbp1+atf6*-cr=38. **E)** Immunofluorescence of type II collagen in the notochord of 28hpf embryos. Simultaneous loss of *creb3l1* and *creb3l2* functions results in collagen retention in notochord cells but does not abolish the glycoprotein-rich ECM sheath (stained by WGA). blue: DAPI; magenta: WGA; yellow: Collagen II. Scale bar: 20um. Embryo orientation is indicated on the bottom left. A: anterior; P: posterior; D: dorsal; V: ventral. **F)** Double *creb3l1+3l2*-cr larvae exhibited shorter body lengths at 5dpf. Scale bar = 1mm.

### Cargo secretion requires different UPR TFs in the notochord and hatching gland

To test the roles of the UPR TFs in notochord and hatching gland differentiation, we performed CRISPR-mediated mutagenesis of the four UPR TFs (*xbp1*, *atf6*, *creb3l1*, and *creb3l2*). Crispant embryos were generated by co-injecting Cas9 protein with guide RNAs at the one-cell stage. As described previously, the >90% high editing rates generated loss-of-function genotypes and did not lead to obvious non-specific developmental defects^55,56^ (**Figure S4A-C**, **table S5**). We additionally raised a stable mutant line for *creb3l1*. The crispant approach (in wildtype or the stable mutant embryos) also allowed testing for redundant functions by generating double, triple, and quadruple loss-of-function embryos, where multiple UPR TFs were simultaneously eliminated.

We found that *xbp1* is required in the hatching gland for successful hatching enzyme (Ctslb) secretion and for hatching, while *creb3l1* and *creb3l2* are redundantly required in the notochord for type II collagen secretion. Specifically, *xbp1* crispants were associated with the absence of Ctslb from the hatching gland cell surface and hatching deficiency, consistent with hatching gland defects observed in zebrafish *xbp1* morphant embryos (**Figure 3C-D**) ^17^. *creb3l1*+*3l2* double crispants exhibited reduced collagen in the notochord sheath, collagen aggregation inside notochord cells, and shortened trunk length, resembling a *creb3l2* hypomorphic mutant (**Figure 3E-F**)^18,21^. Our results additionally showed that, despite being expressed by both the notochord and the hatching gland, *xbp1* is not required in the notochord for collagen secretion, and *atf6* is dispensable in both cell types, contrary to its essential role in mice and Medaka, where *atf6* loss-of-function is embryonic lethal (**Figure 3C-E**)^74–76^. These results demonstrate that UPR TFs are required for secretory function acquisition in the two cell types, but each cell type requires distinct TFs.

### Activation of secretory machinery genes requires different UPR TFs in the notochord and hatching gland

To test the gene regulatory roles of the UPR TFs, we next identified their target genes during notochord and hatching gland differentiation. We define target genes as genes whose expression levels change in response to UPR TF perturbations, which can include both direct and indirect UPR TF targets. scRNA-seq of UPR TF crispants was performed at 12hpf, when UPR TFs and secretory pathway genes, but not cargo proteins, are highly expressed (**Figure 2E**). We analyzed a total of ∼60,000 transcriptomes from 12 experimental conditions, including single, double, triple, and quadruple loss-of-functions (**Figure 4A**, **table S6**). Despite the dozens of genes misregulated in UPR TF crispants (see below), cell type identities were recognizable in the cluster analysis (**Figure 4B**). Target genes for the UPR TFs (*xbp1*, *creb3l1*, *creb3l2*, *atf6*) were inferred by fitting a robust linear regression (RLM) model for each gene in each cell type. This model aims to explain the transcriptional variation of a gene based on the expression levels of the four UPR TFs; a gene is identified as a target if its expression can be effectively explained by UPR TF expression levels (see Method).

**Figure 4.**
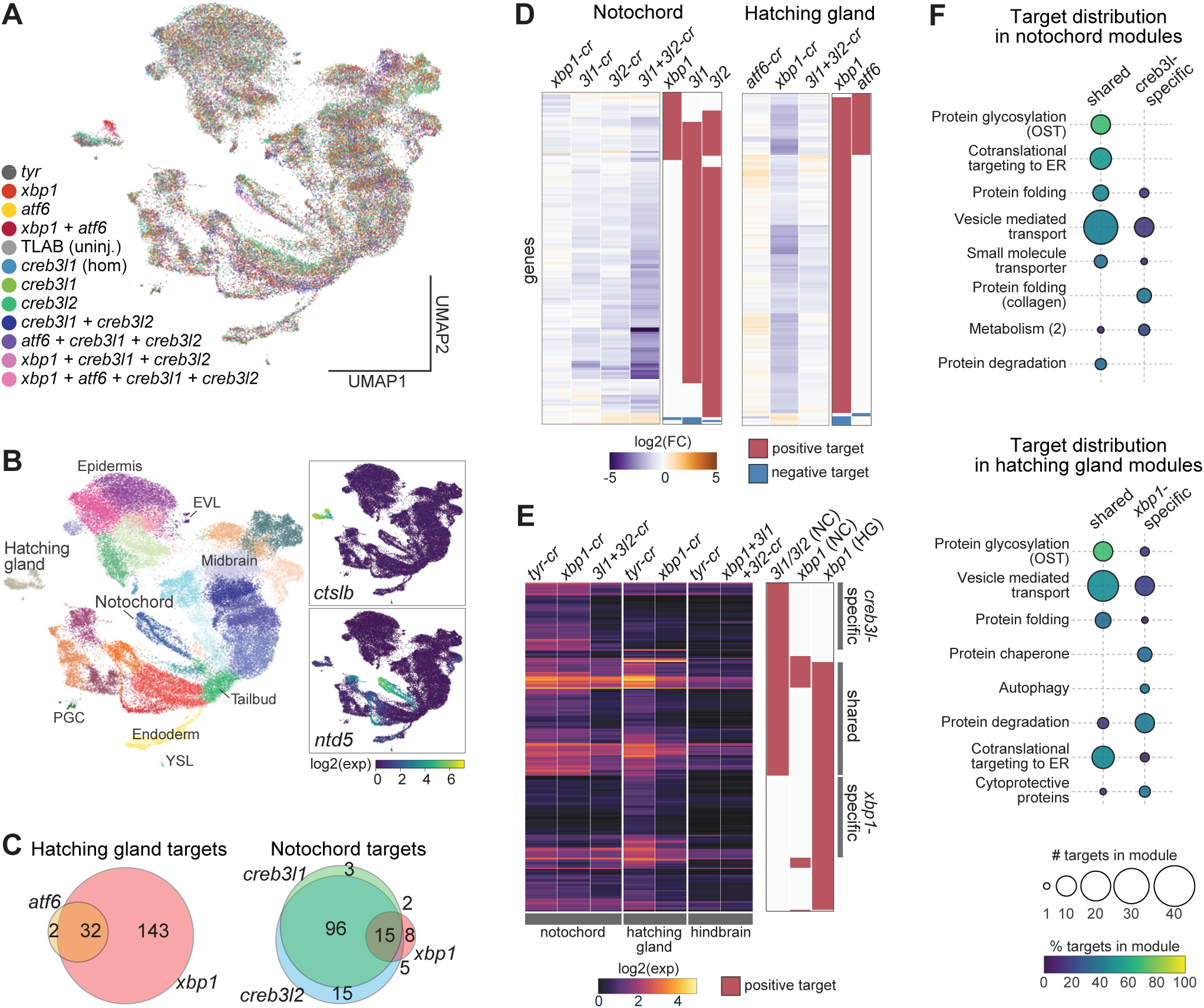
Transcription factors involved in unfolded protein response regulate shared and cell type-specific genes. **A)** UMAP projection of the single-cell transcriptomes colored by sample. “TLAB”: uninjected wild-type. “*creb3l1* (hom)”: *creb3l1* homozygous mutant. Multi-gene loss-of-function conditions involving *creb3l1* are crispants of other genes in *creb3l1* homozygous embryos. All other conditions are crispants of the genes listed. **B)** UMAP colored by cell types (left panel) and by marker genes for the hatching gland (*ctslb*) or notochord (*ntd5*). Cells were clustered using the Leiden algorithm and annotated based on marker gene expression. Several cell types are labeled. Two spatially separated cell populations in the hatching gland cluster exist: one with *xbp1* loss and one without (see color and legend in A). **C)** Venn diagrams of UPR TF targets in the notochord and hatching gland. See also Figure S5A for target gene numbers for each UPR TF in different cell types. **D)** Heatmap of target gene log2 fold changes (UPR TF crispants : *tyr* crispants). Each row is a target gene. Right bar graphs show whether a gene is a target for the TF labeled above. “*3l1*”: *creb3l1*; “*3l2*”: *creb3l2*; “cr”: crispant. **E)** Expression heatmap of *xbp1*, *creb3l1* and *creb3l2* target genes. Rows are targets. Mean expression per cell type is shown. Red bars mark the positive targets for the TF in the cell type labeled on top (“NC”: notochord; “HG”: hatching gland). *creb3l1* and *creb3l2* targets are shown as a union (*3l1/3l2*). Gray bars on the right mark the three classes of targets as defined in the main text. **F)** Distribution of the three classes of target genes in the previously identified hatching gland and notochord gene modules. Modules are ordered by their target gene percentages, i.e. the percentage of genes in a module that are targets. Only the top modules in each cell type are shown.

We found that *xbp1* positively regulates 175 genes in the hatching gland but only 30 genes in the notochord (**Figure 4C, Figure S5A, Table S7**). Only a handful of these targets (e.g. genes involved in translocation of nascent proteins into ER) were identified in non-hatching gland cells (e.g. epidermis), where *xbp1* is expressed at a much lower level compared to the hatching gland (**Figure 3A**, **Figure S5B-C**). *creb3l1* and *creb3l2* each regulate over 110 genes in the notochord but no genes in the hatching gland (**Figure 4C, Table S7**). They exhibited highly redundant regulatory functions in the notochord, as their target genes were largely identical and showed strong down-regulation only in the *creb3l1*+*3l2* double crispants (**Figure 4C-D**). *atf6* regulated few genes in either cell type (**Figure 4C, Figure S5A, Table S7**). Few or no negatively regulated targets were identified for any UPR TF (**Figure S5A**). These results are highly consistent with our phenotyping experiments (**Figure 3C-F**). Additionally, loss of UPR TFs reduced target gene expression to basal levels comparable to cells with non-secretory fates, arguing that UPR TFs are the major activators of these secretory pathway genes in the differentiating notochord and hatching gland (**Figure 4E**).

Altogether, these results indicate that UPR TFs are primarily active in the axial mesoderm during embryogenesis, where they induce secretory pathway genes during notochord and hatching gland differentiation prior to large-scale cargo production. Further, distinct UPR TFs activate secretory pathway genes in each cell type — namely, *xbp1* in the hatching gland, and *creb3l1* and *creb3l2* redundantly in the notochord.

### *creb3l1*/*3l2* and *xbp1* regulate ECM secretion and gland-like secretion programs, respectively

To determine the functions controlled by each UPR TF, we inspected the functional annotations of their target genes. ∼85% of the UPR TF target genes were included in our previously defined gene modules. These targets were strongly enriched in modules associated with secretory processes in both cell types, such as nascent protein cotranslational targeting to the ER, protein glycosylation by oligosaccharyltransferase (OST), protein folding, and vesicle mediated transport (**Figure 4F**). In contrast, UPR TFs did not activate genes associated with non-secretory functions, such as cell specification and tissue patterning.

To assess functional associations with distinct modes of regulation, we classified the target genes into three classes: shared targets (regulated by *xbp1* in the hatching gland AND *creb3l1*/*3l2* in the notochord), *creb3l*-specific targets (higher expression in the notochord and targets of *creb3l1*/*3l2* but NOT *xbp1)*, and *xbp1*-specific targets (higher expression in the hatching gland and targets of *xbp1* but NOT *creb3l1*/*3l2*) (**Figure 4E-F, Table S7**). 80 genes were shared targets, and they contribute to several common secretory steps, including co-translational targeting to ER (e.g. *sec61g*, *srpr*, *spcs1*), protein folding (e.g. *p4hb*, *pdia6*, *fkbp9*), vesicle mediated transport (e.g. *copa*, *kdelr3*, *sec22bb*), and OST-mediated glycosylation (e.g. *ost4*, *rpn1*, *ddost*) (**Figure 4F**). We refer to these targets as a ‘general secretion’ program.

The 58 *creb3l*-specific targets include enzymes specialized in collagen folding (e.g. *crtap*, *p3h1*)^57^ and vesicle-mediated transport components that specifically accommodate large ECM proteins (e.g. *sec23A*, *sec13*)^58^ (**Figure 4F**). We refer to the *creb3l*-specific targets as an ‘ECM secretion’ program. Although the *creb3l1/3l2* regulated secretion program appears to be tailored for secreting ECM components, ECM components such as collagens are regulated independently, as their expression was largely unaffected in the *creb3l1*+*3l2* crispants (**Figure S5D**).

The 47 *xbp1*-specific targets include many ER chaperones (e.g. *dnajc3a*, *sil1*) and ER-associated protein degradation (ERAD) machinery (e.g. *edem2*, *syvn1*) (**Figure 4E**)^59^. We refer to the *xbp1*-specific targets as the ‘gland-like secretion’ program. The hatching gland may require these additional chaperones and degradation machinery to maintain ER homeostasis during this tissue’s extremely high load of secretory protein production (**Figure 2E**). Analogous to collagens in the notochord, the hatching enzymes are not regulated by the hatching gland UPR TF *xbp1* (**Figure S5D**). Therefore, UPR TFs activate secretory machineries tailored to specific cargos, while secretory cargos themselves are regulated independently.

In summary, common secretory pathway genes are activated by distinct UPR TFs in the two cell types — by *xbp1* in the hatching gland and redundantly by *creb3l1* and *creb3l2* in the notochord. In addition, each UPR TF activates a more specialized program tailored to the secretory cargo produced by each cell type – namely, *xbp1* activates a gland-like secretion program, while *creb3l1/3l2* redundantly activates an ECM secretion program.

### *creb3l1/3l2* are sufficient to fully activate the ECM secretion program, while *xbp1* only partially activates the gland-like secretion program

Since *xbp1* and *creb3l1*/*3l2* each regulate a set of cell-type-specific genes in the hatching gland and notochord, respectively, we wondered how this specificity is conferred. In the simplest model, *xbp1* and *creb3l1*/*3l2* are sufficient to specifically regulate the gland-like and ECM secretion programs, respectively. In this model, the ECM secretion program is restricted to the notochord because *creb3l1* and *creb3l2* are only expressed in the notochord, and mis-expression of *creb3l1*/*3l2* in another cell type would activate the ECM secretion program there as well. Alternatively, the UPR TFs might cooperate with cell-type-specific TFs to confer specificity, in which case the mis-expression of the UPR TFs alone would not be sufficient to activate these specific secretion programs in other cell types.

To distinguish between these and other models, we globally mis-expressed each UPR TF by injecting mRNAs encoding their activated forms (**Figure 5A**). We then performed scRNA-seq on the resulting embryos at 8hpf, when 1) a sizable population of transcriptionally distinct notochord and hatching gland cells are detectable (**Figure 1B**), and 2) the endogenous UPR TFs and secretory pathway genes are not yet highly expressed (except for *xbp1*) (**Figure 1D**). Since *xbp1* can be activated by UPR TFs including *creb3l1*, *creb3l2*, and *atf6*, we mis-expressed the UPR TFs in *xbp1* crispants to reduce potential indirect effects (**Figure S6A**). Single-cell transcriptomes were obtained across 5 mis-expression (“me”) conditions, including an mCherry control, the activated forms of *xbp1* (*xbp1-s*), *creb3l1* (*creb3l1-N*), *creb3l2* (*creb3l2-N*), and *atf6* (*atf6-N*) (**Figure 5B**).

**Figure 5.**
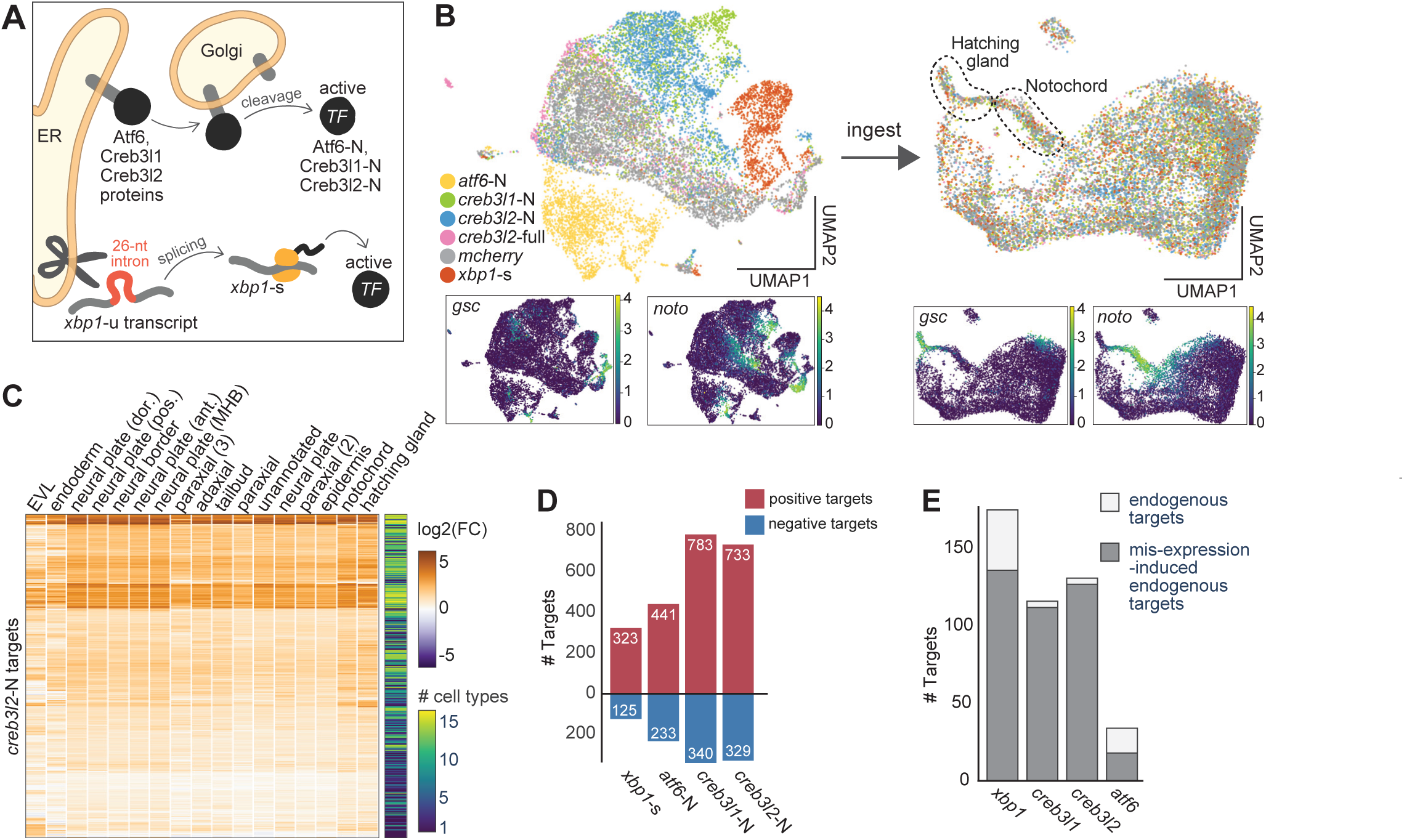
*creb3l1/3l2* are sufficient to activate the extracellular matrix secretion program. **A)** Schematic of UPR TF activation upon ER stress. Unspliced *xbp1* transcript (*xbp1*-u) is spliced to produce the *xbp1-s* isoform, which encodes the active TF. Atf6, Creb3l1 and Creb3l2 proteins are first translocated to the Golgi, then cleaved to release their cytoplasmic domains (N-termini) (Atf6-N, Creb3l1-N, Creb3l2-N), which then function as active TFs. **B)** Cell type identification for the UPR TF mis-expression scRNA-seq data. UMAP projection before (left) and after (right) dataset integration by the ingest algorithm are shown. UMAPs are colored by samples (upper) or marker gene expression (lower). *gsc* and *noto* are hatching gland progenitor and notochord markers, respectively. Before ingest integration, transcriptomes from different conditions separated on the UMAP due to the global transcriptional profile shifts induced by UPR TF mis-expression. Ingest mapped transcriptomes from different samples to the control data (mCherry) manifold without altering the original read counts. This allowed cell type labels to be transferred from the control to all samples. **C)** Heatmap of the log2 fold changes of *creb3l2-*N positive targets (rows) by cell types (columns). The 1436 genes detected as positive targets in at least one cell type are shown. Right side bar shows the number of cell types in which each gene is identified as a target. *creb3l2-*N shows consistent gene induction activities across cell types, except for in EVL and endoderm. See Figure S6B for similar plots for all other mis-expressed UPR TFs. **D)** Number of mis-expression regulated targets by UPR TFs. Genes identified as a target in only one cell type were excluded. Negative bars represent negative targets, i.e. genes downregulated in mis-expression samples. **E)** Mis-expression of active UPR TFs induced many of their endogenous targets. Only positive targets are shown.

We inferred target genes of each UPR TF within individual cell types using RLM, similarly to our analysis of the crispant data (Methods). We observed that each UPR TF activates a highly consistent set of genes across most cell types in the embryo (**Figure 5C, Figure S6B**). To identify robustly induced genes and reduce false positives, we defined “induced targets” for each UPR TF as the genes identified as positive targets in at least two cell types (**Figure 5D, Table S7**). The induced targets included many that were down regulated in the crispants. Specifically, *creb3l1-N* and *creb3l2-N* were sufficient to induce almost all their endogenous targets, while *xbp1-s* and *atf6-N* activated a smaller subset of their respective endogenous targets (**Figure 5E, Figure S6C).**

We next asked whether the general, ECM, and gland-like secretion programs were activated by the mis-expressed UPR TFs. Our results showed that the general secretion program was fully activated by either *creb3l1-N*, *creb3l2-N*, or *xbp1-s*, but not *atf6-N*. The ECM secretion program was fully activated by *creb3l1-N* or *creb3l2-N*, but not by *xbp1-s* or *atf6-N*. Finally, the gland-like secretion program was partially activated by *xbp1-s*, and a smaller portion of the program was activated by *creb3l1-N, creb3l2-N,* and *atf6-N* (**Figure 6A, Figure S6D**).

**Figure 6.**
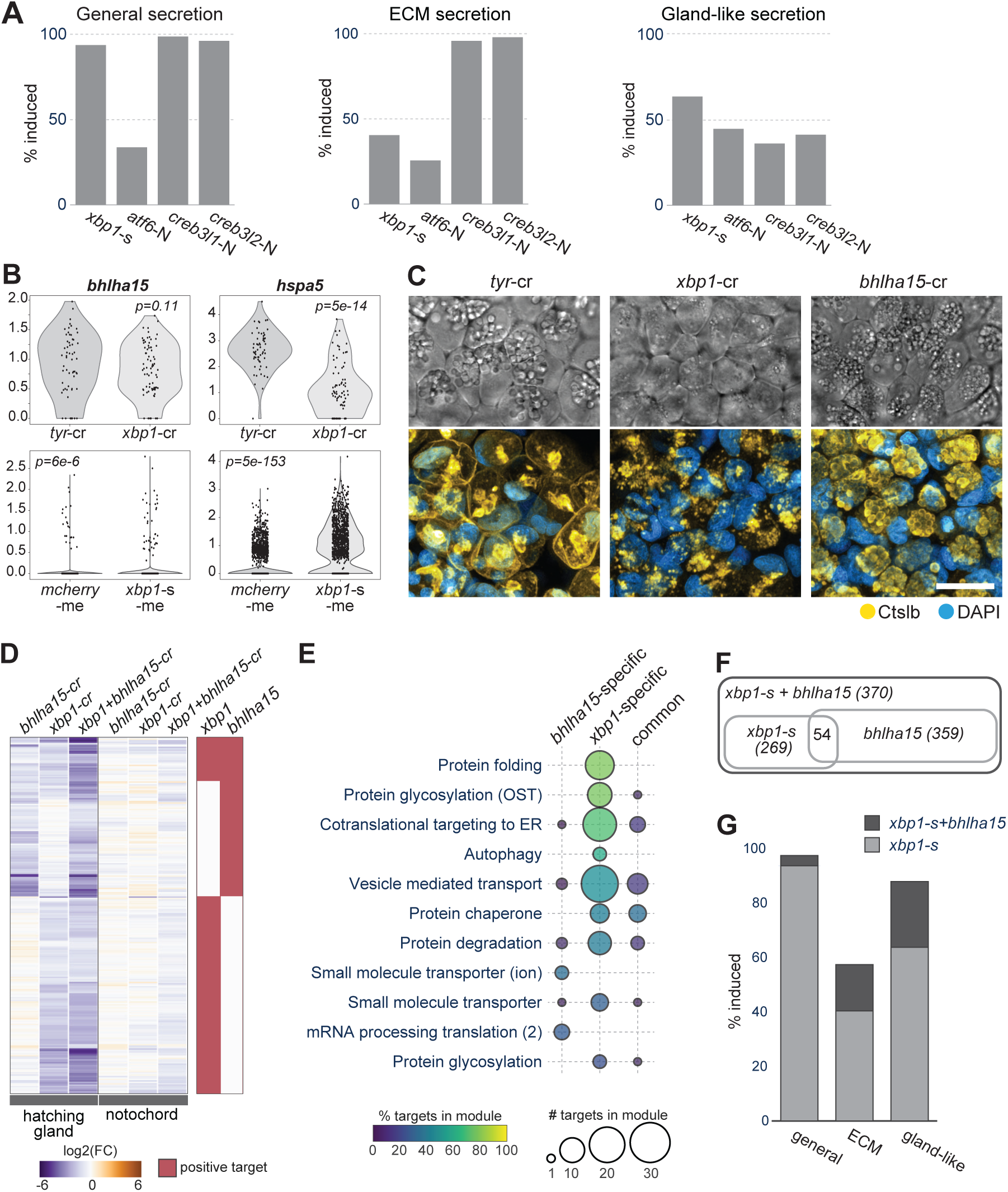
*xbp1* partners with *bhlha15* to activate the gland-like secretion program. **A)** Percentage of genes induced by UPR TF misexpression in the general, ECM, and gland-like secretion programs, which include 80, 47, and 58, respectively. **B)** *bhlha15* is not an endogenous or induced target of *xbp1*. *bhlha15* expression did not show significant change in *xbp1* loss- (upper) or gain- (lower) of function embryos. *hspa5* is shown as an example *xbp1* target. P-value in each plot is resulted from a t-test of the expressions between two samples. **C)** *bhlha15* crispants exhibit similar but less severe hatching gland defects compared to *xbp1* crispants at 28hpf. scale bar = 20um. **D)** A subset of *bhlha15* targets are shared with *xbp1*. Heatmap shows log2 fold change of target genes (rows). Red bars indicate whether each gene is a target in the hatching gland for the TF labeled on top. “cr”: crispant. **E)** Distribution of *xbp1* and *bhlha15* endogenous targets in hatching gland gene modules. **F)** Overlap between targets induced by mis-expressing *xbp1-s*, *bhlha15*, and *xbp1-s+bhlha15*. **G)** *xbp1*-s+*bhlha15* co-mis-expression induced additional genes in the general, ECM, and gland-like secretion programs compared to *xbp1*-s mis-expression alone.

These results indicate that *xbp1-s*, *creb3l1-N* and *creb3l2-N* can act as the master regulators of the general secretion program. In addition, *creb3l1-N* and *creb3l2-N* can act as the master regulators of the ECM secretion program, whereas full induction of the gland-like secretion program cannot be achieved by *xbp1-s* activity alone.

### *xbp1* requires *bhlha15* to fully activate the gland-like secretion program

Many gland-like secretory genes were downregulated in the hatching gland in *xbp1* loss-of-function individuals, but not activated upon *xbp1-s* mis-expression. We hypothesized that *xbp1* may require hatching gland specific transcription factors to cooperatively activate the gland-like secretory program. To identify cooperative factors of *xbp1*, we used a candidate approach and considered TFs enriched along the hatching gland trajectory whose expression begins before the expression of secretory pathway genes. Out of the 34 candidate TFs, 25 were enriched in the hatching gland and not the notochord. Among these, *bhlha15*/*mist1* from the transcription regulation module was previously described as a potential *xbp1* target gene: several studies in pancreatic cell differentiation and drug induced UPR have suggested that *xbp1* activates *bhlha15*, which in turn cooperates with *xbp1* to activate additional secretory pathway genes^43,60,61^. Although *bhlha15* was not identified as an *xbp1* target from either our crispant or mis-expression experiments (**Figure 6B**), we hypothesized that it might still cooperate with *xbp1* in the hatching gland.

To test the role of *bhlha15*, we generated crispants. *bhlha15* crispant embryos exhibited low hatching rate (**Figure S7A**) and hatching enzyme (Ctslb) secretion deficiency similar to *xbp1* crispants (**Figure 6C**). In addition, both *xbp1* and *bhlha15* crispants formed smaller secretion granules in the hatching gland cells compared to the control, but with a less severe phenotype in *bhlha15* crispants (**Figure 6C**). These observations establish the requirement of *bhlha15* in hatching gland secretion.

To test which hatching gland genes are regulated by *bhlha15*, we performed scRNA-seq on *bhlha15* crispants, as well as on *bhlha15*+*xbp1* double crispants. Target gene analysis by RLM revealed that *bhlha15* regulates a subset (18.4%) of *xbp1* endogenous targets, as well as a larger set of genes not regulated by *xbp1* (**Figure 6D, Table S7**). Most of the *bhlha15*-specific targets were members of gene modules involved in secretion, but no particular module showed high representation of *bhlha15*-specific targets (**Figure 6E**).

To investigate whether *bhlha15* and *xbp1* together activate the gland-like secretion program, we conducted scRNA-seq at 8hpf for embryos mis-expressing *xbp1-s*, *bhlha15*, or *xbp1-s*+*bhlha15* (**Figure 6F**, **Figure S7B**). While mis-expression of *xbp1-s* or *bhlha15* alone induced a smaller subset of the gland-like secretion program, *xbp1-s*+*bhlha15* co-mis-expression activated almost all the gland-like secretion genes but only half of ECM secretion program genes (**Figure 6G, Table S7**). Taken together, these results indicate that the combination of *bhlha15* and *xbp1* is sufficient to activate the gland-like secretion program in diverse embryonic cell types.

## Discussion

In this study, we comprehensively reconstructed the cellular remodeling processes during notochord and hatching gland differentiation and dissected the regulatory logic of secretory machinery remodeling. Our study introduces a generalizable approach for the genome-wide exploration of cellular differentiation and proposes a regulatory model for cell-type specific and shared differentiation programs (**Figure 7**).

**Figure 7.**
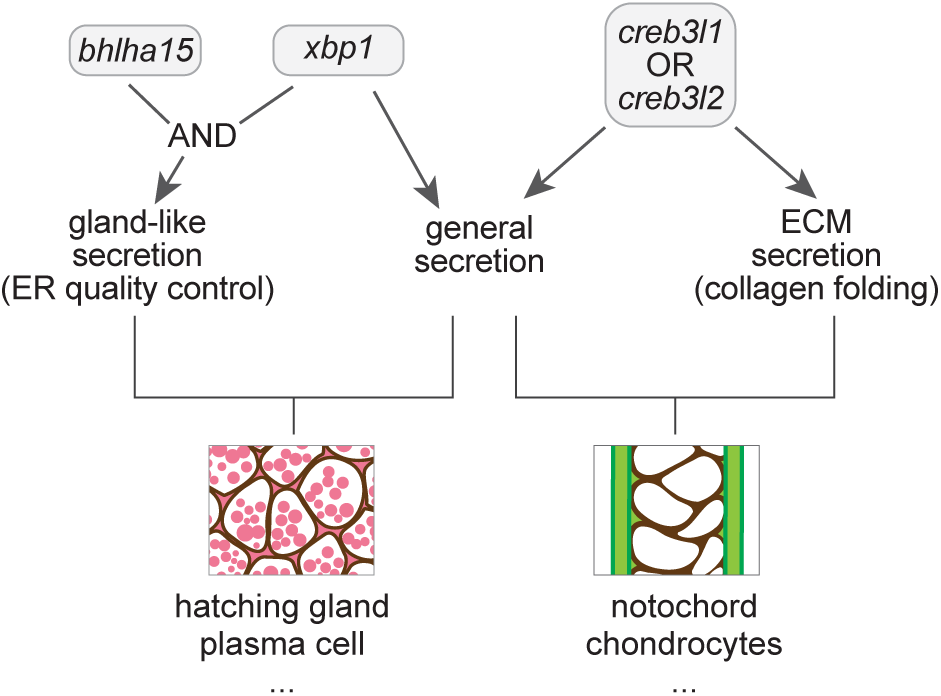
Model for the activation of shared and specific secretion programs by unfolded protein response transcription factors.

First, we established a data mining and computational approach (MIMIR) that leverages information from both time course scRNA-seq data and functional annotation databases to define gene modules during cell type differentiation. Our approach extends previous approaches that have mostly relied on gene ontology (GO) and pathway enrichment (e.g. KEGG). These approaches aim to identify the biological processes or pathways statistically enriched in a set of genes of interest^62–66^. While suitable for identifying the major biological processes, such methods have several limitations: 1) functional annotations are not available for all genes in each database, omitting many genes from the analysis; 2) statistically significant processes in a tissue often cover only a subset of all functionally annotated genes, leaving even more genes out of analysis; and 3) many genes are associated with multiple enriched processes, making their context-specific functions ambiguous. To recover functions for more genes and to resolve ambiguous gene functions, we extended the gene ontology and pathway enrichment analysis by 1) using a combination of databases, namely GO, STRING, Reactome, and InterPro^24–27^, 2) incorporating expression dynamics from transcriptional trajectories, and 3) assigning genes into distinct, non-overlapping functional modules. This approach combined with manual curation assigned a high percentage of all input genes (∼95%) into specific functional modules. Moreover, distinct modules covering most enriched genes (∼90%) could be identified with automated clustering alone. MIMIR should be applicable to any differentiation process for which time course RNA-seq data is available.

Second, the identified gene modules offer a comprehensive view of the cellular processes in notochord and hatching gland differentiation. These modules 1) recovered known cellular processes, such as specification, cytoskeletal remodeling, and vesicle-mediated transport, 2) identified newly associated cellular processes in these tissues, such as mitochondria function and tRNA processing, 3) associated previously uncharacterized genes to known biological processes, such as the new notochord vacuolation gene – *atp6v0e1* (**Figure S2**), 4) demonstrated that different cell types can use distinct genes to control similar processes (**Figure 2C**), and 5) revealed a preparatory organization of secretory function development, with UPR transducers and secretory machineries expressed prior to cargo protein production. These results suggest that module analysis facilitates the characterization of cellular remodeling and uncovers previously neglected processes and genes. Extension of module analysis to other cell types and organisms promises to advance our understanding of differentiation.

Third, our study illustrates the power of combining scRNA-seq with loss- and gain-of-funtion manipulations to phenotype embryos through *in silico* cell sorting^56,67^. We show that the F0 crispants exhibit loss-of-function phenotypes with little or no unspecific developmental defects on both morphological and transcriptional levels (**Figure S4**). The crispant approach enabled us to analyze complex genetic perturbations, including quadruple loss-of-functions and mis-expression in mutant backgrounds, to determine necessity, sufficiency, and redundancy in loss- and gain-of-function experiments. Many of these combinations would have been challenging to construct and analyze with stable mutants or transgenics. For example, the double loss-of-function embryos revealed the redundancy of *creb3l1/3l2* in target gene regulation in the notochord. Analogously, the gain-of-function embryos revealed the sufficiency (*creb3l1/3l2*) and co-dependence (*xbp1* + *bhlha15*) of UPR TFs in non-axial mesoderm cells. Thus, the combination of CRISPR-mediated mutagenesis, RNA-mediated mis-expression, scRNA-seq and *in silico* cell sorting can provide organism- and genome-wide insights into complex genetic interactions and gene regulatory networks.

Fourth, we shed light on the regulatory logic of general and cell-type-specific secretory functions by UPR TFs. *xbp1*, *creb3l1*, and *creb3l2* each activate a general secretion program; *creb3l1* and *creb3l2* additionally activate an ECM-secretion program, while *xbp1*, together with *bhlha15*, regulates a gland-like secretion program (**Figure 7**). Our study extends previous studies of the roles of UPR TFs in ECM-secreting cells (*e.g.* chondrocytes) and gland-like secretory cells (*e.g.* pancreatic acinar cells) by identifying target genes on a genome-wide scale and assessing the specificity and interchangeability of these TFs^17,18,21,52,68,69^. Together, these studies establish that *creb3l1/3l2* and *xbp1* activate a shared, general secretion program that includes genes involved in synthesizing and transporting secretory proteins. However, these UPR TFs remain non-interchangeable due to their specificity for ECM or gland-like secretion programs, respectively. Our mis-expression and loss-of-function analyses suggest that UPR TFs can serve as key regulators, or master regulators, in different contexts to generate specialized secretory functions: *creb3l1*/*3l2* confer ECM secretion functions and induce genes involved in collagen folding and transport; *xbp1* and *bhlha15* together activate the gland-like secretory program and induce genes involved in ER quality control.

In addition, our results reveal an intriguing phenomenon where target genes of UPR TFs are expressed even before the accumulation of high levels of protein cargos. This observation suggests an anticipatory response to cellular stress that would be caused by large-scale cargo production. This anticipatory activity of UPR TFs could be regulated at both transcriptional and post-transcriptional levels. Transcriptionally, certain UPR TFs are developmentally programmed; for instance, *xbp1* expression in zebrafish hatching gland progenitors is directly activated at the onset of gastrulation by Nodal signaling, and in mouse chondrocytes, *creb3l2* can be activated by Sox9^17,70^. Post-transcriptionally, these TFs often require activation, as seen with *xbp1* undergoing splicing in zebrafish embryos, and reminiscent of the early *xbp1*-s detection in B-cell differentiation prior to substantial immunoglobulin production^71^. Furthermore, we found that the cleaved form of Creb3l2 is more potent than its unprocessed version, indicating the necessity of post-translational activation for full functionality (**Figure 5B**, **Figure S6C**)^17,19^. Identifying the triggers for this post-transcriptional activation remains an open question. Potential mechanisms include the production of misfolded proteins before classical cargo accumulation, higher basal processing rates in embryos, changes in ER membrane lipid composition^72^, or other undiscovered pathways leading to the anticipatory activation of UPR TFs.

More generally, the regulatory roles of *creb3l1/3l2*, *xbp1*, and *bhlha15* provide new insights into the logic of gene regulation during differentiation (**Figure 7**). First, the activation of the general secretion program by *creb3l1*/*3l2* or *xbp1* in different cell types is reminiscent of the use of distinct but related TFs in activating shared neurotransmitter synthesis pathways in different neurons^47,48^. This process of phenotypic convergence allows these different TFs to activate a shared gene expression program in different cells. Second, UPR TFs also have features of master regulators of specialized differentiation programs. Akin to factors that can activate the full gene expression program for ciliogenesis, *creb3l1*/*3l2* is a master regulator of the ECM secretion program and is sufficient to activate the associated genes in different cell types^2,46^. Third, the activation of the gland-like secretory program by *xbp1* and *bhlha15* illustrates a combinatorial strategy: the need for cell-type specific cofactors to confer cell type specificity in activating a differentiation program, as observed in the differentiation of distinct muscle types^73^. Thus, UPR TFs use a unique combination of phenotypic convergence, master regulation, and combinatorial regulation to confer shared and cargo-specific secretory features and are a fascinating paradigm to dissect the regulatory logic of differentiation.

### Limitations of the study

Our study has three main limitations. First, although the automated module identification approach identified distinct modules that cover most enriched genes, manual curation was necessary to improve the accuracy of module memberships, to annotate the modules, and to assign additional enriched genes to modules. We expect that future integration of additional sources of information (e.g. co-expression data under different conditions; co-expression data across organisms; ATAC-seq data; morphological data) will reduce curation and further enhance module identification. Second, our target gene analysis is based on scRNA-seq data and therefore cannot distinguish between direct and indirect targets. Future studies that detect UPR TF binding sites with cell-type-resolution will help address this limitation. Finally, our analysis suggests an anticipatory role of the UPR TFs: UPR targets that expands secretory capacities are already expressed before high levels of cargo are present. Future studies will be needed to identify the factors that trigger the early post-transcriptional activation of UPR TFs in differentiating secretory cells.

## Acknowledgments

We thank K. Shekhar, D. Ron, K.C. Sadler, M.C. Tobias, M. Colomer-Rosell, J. Navajas Acedo, O. Mayseless, A.L.A. Nichols, Y. Wan, and E.A. Bayer for feedback on the manuscript. We thank the Harvard Center for Biological Imaging and Bauer Core Facility, the Genomics Facility Basel, and the Biozentrum Imaging Core Facility for experimental support. We thank Harvard FAS Research Computing and Biozentrum SciCore for providing computational resources. This work was funded by ERC Advanced grant 834788 (AFS), the Allen Discovery Center for Cell Lineage Tracing (AFS), NIH K99HD091291 (JAF) and ZIAHD008997 (JAF).

## Author contributions

Conceptualization, Y.W., J.A.F., and A.F.S., J.L.; Methodology, Y.W., J.A.F., and A.F.S.; Formal Analysis, Y.W., J.A.F., and J.L.; Investigation, Y.W., J.A.F., J.L., L.Y.D., and J.L.W.; Writing – Original Draft, Y.W., J.A.F., L.Y.D., and A.F.S.; Funding Acquisition, A.F.S. and J.A.F.

## Declaration of Interests

The authors declare no competing interests.

## Inclusion and Diversity

We support inclusive, diverse, and equitable conduct of research. This study was performed by authors of multiple races, genders, and sexual orientations.

## Methods

### Resource Availability

#### Lead Contact

Requests for further information, resources, and reagents should be directed to the lead contact Alexander F. Schier.

#### Materials availability

Plasmids generated in this study will be shared by the lead contact upon request. Mutant zebrafish lines generated in this study will be shared by the lead contact upon reasonable request (subject to import/export restrictions).

#### Data and Code Availability

Sequencing data is available as FASTQs and UMI count tables under NCBI GEO accession GSE252032. Code for gene module identification will be available at https://github.com/YiqunW/MIMIR.

### Method Details

#### Reconstruction of Axial Mesoderm Transcriptional Trajectories

Single cell transcriptomes from 3.3-12hpf were obtained from^1^. Additional single cell transcriptomes from 12hpf (10-somite) to 24hpf were obtained from^2^. Single cell trajectories were previously reconstructed using the dataset spanning 3.3-12 hpf, which included the trajectories for notochord and the hatching gland (also known as the prechordal plate in the previous study)^1^. Cells along these trajectories were isolated and combined with the notochord and hatching gland cells extracted from the dataset that covers later developmental stages. Notochord and hatching gland cells were identified from the second dataset by clustering data from each timepoint and inspecting marker gene expressions (*noto* and *ntd5* for the notochord; *ctsl1b* and *cd63* for the hatching gland). Clustering was performed using variable genes identified as previously described^3^. Briefly, a null model (negative binomial distribution) was fitted to our the 12-24hpf data to explain the expected technical variation for each gene, given its expression level. Genes were considered to be ‘variable’ if their coefficient of variation (CV) sufficiently exceeded the null model (CV >= 0.5). A community detection algorithm was used to cluster the cells based on their variable gene expression (the function graphClustering() from the URD package was called with argument method=c(“Infomap”)^1^).

The extended transcriptional trajectories for axial mesoderm from 3.3hpf to 24hpf were reconstructed from the combined dataset using the URD package^1^. Cells from the earliest stage (3.3hpf) were used as the ‘root’, and notochord and hatching gland cells from the last stage (24hpf) were used as the two ‘tips’ during pseudotime calculation and trajectory inference.

#### Identification of Enriched Genes

Identification of enriched genes from 3.3 to 12hpf:

We first used our previously constructed transcriptional trajectories for the whole embryo^1^ to identify enriched genes up to 12 hpf in the notochord and the hatching gland. The pseudotime line was first divided into 15 equal segments. If a segment in a trajectory contains fewer than 60 cells, we fused it with its adjacent segment(s) until the number of cells in each segment is greater than 60. When multiple consecutive segments had fewer than 60 cells, we fused them in a way so that the final number of cells in different segments are as even as possible. 9 and 8 final segments were defined along the notochord and the hatching gland trajectories, respectively.

For each cell type, we searched for marker genes in each pseudotime segment separately. For instance, for the notochord, we compared the notochord cells in a pseudotime segment with all cells that are not axial mesoderm (not notochord or hatching gland) in the same pseudotime window. We used a precision-recall curve (AUCPR) to identify genes that can separate the two groups of cells well in the segment^3^. A gene is identified as a notochord marker gene in a segment if it meets all following criteria: 1) it is detected in at least 10% of the cells in the notochord segment, 2) its AUCPR value is at least 2 times the value expected from randomized data, and 3) the log fold change of average expression between the notochord and the non-axial mesoderm cells must be greater than 0.3. Notochord enriched genes were then defined as those that are either a marker gene for the last psuedotime segment, or a marker gene for at least two segments. Enriched genes for the hatching gland were defined the same way.

Identification of enriched genes from 12 to 24hpf:

Our single cell transcriptomes from 12-24hpf covers only the anterior part of the embryo^2^. Since we did not reconstruct trajectories for non-axial-mesodermal cell types in the dataset, pseudotime labels were only available for the axial mesoderm cells. Instead of comparing axial mesoderm with non-axial-mesoderm cells by psudotime segments, we compared the two groups of cells by developmental stages. For each axial mesoderm cell type at each stage, we defined marker genes similarly as we did for the previous dataset. Namely, a precision-recall curve was fit to access how well each gene can differentiate the axial mesoderm cells from the non-axial mesoderm cells at each developmental stage. A marker gene at each developmental stage needs to be detected in at least 10% of the cells in the cell type it marks, has an AUCPR value is at least 7.5 times the value expected from randomized data (a higher value was used due to the fewer cells detected in this dataset), and an expression log fold change greater than 0.3 compared to the non-axial mesoderm cells. Genes identified as a marker gene at any developmental stage are considered an enriched gene.

The final list of enriched genes for each cell type is the union of the enriched genes found from the two datasets.

#### Gene Ontology Enrichment Analysis

Gene Ontology (GO) term enrichment tests were done using Fisher’s exact test. Genes enriched in each of the 25 cell types in the 3-24hpf trajectory data were identified and combined to serve as the background genes.

#### Identification of Genes Modules

Calculating gene expression similarities:

Gene modules are identified based on genes expression similarities as well as their functional similarities. To calculate gene expression similarities, we used the data matrix of gene expression by pseudotime for each cell type. We excluded the earliest pseudotime segment corresponded to cells that are not yet specified as axial mesoderm. These cells were identified as the ones before the branchpoint that trifurcated into the ectoderm, axial mesoderm, and other mesendoderm in the reconstructed trajectories^1^. Expression similarities between each pair of genes were then calculated with the remaining cells for each cell type. Expression similarities were calculated as (1 - expression_dist), where expression_dist is the expression distance scaled to range between 0 and 0.95. We tested a number of expression distance measures, including Euclidean distance, Cosine distance, Canberra distance, and Jensen-Shannon divergence.

Calculating functional similarities:

To obtain functional similarities between genes, we downloaded the protein interaction score table from the STRING database^4–6^. We used the full protein links table from STRING v11.0 for zebrafish (organism code 7955). STRING uses multiple evidence sources for calculating protein-protein-interaction (PPI) scores, including neighborhood, neighborhood transferred, fusion, cooccurence, homology, coexpression, coexpression transferred, experiments, experiments transferred, database, database transferred, textmining, textmining transferred. For explanations of the evidence source, see^4^. The “database” evidence from STRING is derived from a variety of functional and pathway annotation databases, but which databases and how they are combined into scores is not disclosed. To control for which databases to include, we excluded the “database” and “database transferred” evidence sources from STRING and added additional self-calculated evidence sources using several databases, including GO, Interpro, Reactome, and HGNC. Similarities between pairs of genes from each functional annotation database were calculated based on the semantic similarities among the annotated terms.

Functional databases often contain annotation terms that are hierarchically organized, and the idea behind semantic similarity between annotation terms is that annotations for similar functions should have a close relationship in the hierarchical structure. Functional similarities between genes can then be defined by the semantic similarities between the terms they are annotated with. We adopted algorithms in the R package GOSemSim^7,8^ to calculate the semantic similarities between genes based on each functional annotation database, with parameters ‘measure = "Wang", combine = "BMA"’.

PPI scores from STRING database and the self-calculated semantic similarity scores were then combined to give a single functional similarity score between each pair of genes. Each score is in the range between 0 and 1. Scores were combined according to instructions available on STRING database webpage, namely: 𝑆*_total_* = 1 − (1 − 𝑝) ∏*_i_* [1 − (𝑆_𝑖_ − 𝑝)/(1 − 𝑝)], where *S*_i_ denotes the score from evidence source *i*, and *p=0.41* is the fixed prior value.

Gene network clustering and module curation:

The expression similarity scores, 𝑆_𝑒𝑥𝑝_, and the functional similarity scores, 𝑆_𝑓𝑢𝑛_ , are both bounded between 0 and 1. We tested a few methods of combining the two scores, including the ‘OR’ logic: 𝑆_𝑓𝑖𝑛𝑎𝑙_ = 1 − (1 − 𝑆_𝑒𝑥𝑝_)(1 − 𝑆_𝑓𝑢𝑛_ ), which associates two genes strongly if either expression or functional similarity is high, the ‘AND’ logic: 𝑆_𝑓𝑖𝑛𝑎𝑙_ = 𝑆_𝑒𝑥𝑝_𝑆_𝑓𝑢𝑛_ , which associates two genes only if both their expression and functional similarities are high, and the simple ‘+’ method: 𝑆_𝑓𝑖𝑛𝑎𝑙_ = 𝑆_𝑒𝑥𝑝_ + 𝑆_𝑓𝑢𝑛_ . We built a network of enriched genes connected by the final similarity scores for each cell type, and performed gene clustering using ‘leiden’, ‘louvain’, or ‘infomap’ algorithms from the igraph package^9^. In general, Leiden was able to better partition the densely connected network into biologically meaningful clusters, while other algorithms found relatively few clusters, with most genes encoding secretion pathway components and ECM proteins all clustered together. Combinations of different clustering methods and similarity scores were applied to identify clusters out of a test gene set that included 300 notochord enriched genes. The resulting clusters were compared against the manually curated modules using AMI (adjusted mutual information), computed with the AMI() function from the aricode package. The combination of scores and methods that gave the highest AMI was used in gene module identification. Specifically, expression similarity based on Jensen-Shannon divergence and functional similarity aggregated from STRING (excluding “database” and “database transferred”), GO, Interpro, and Reactome were used. The two modes of similarities were multiplied to give the final similarity score (𝑆_𝑓𝑖𝑛𝑎𝑙_ = 𝑆_𝑒𝑥𝑝_𝑆_𝑓𝑢𝑛_ ), which were then provided as the input to Leiden clustering (with parameters ‘resolution_parameter=4, partition_type = “RBConfigurationVertexPartition”’ in the igraph leiden function).

After obtaining the gene clusters, we performed manual curation to annotate and refine the clusters to generate the final modules. We used the functional annotations enriched in each cluster as the basis for module annotation and curation. If most genes in a cluster is annotated with the same functional term, the cluster will be annotated with that term, and genes in the cluster not annotated with the same term will be inspected and either kept, reassigned, or become a new cluster. Genes whose expression profile clearly deviates from other cluster members will be reassessed as well. If genes in a cluster are annotated with diverse terms, their expression and functions will be reexamined to either identify potential unifying functions or split the cluster. During this curation process, genes’ general functions and their functional relevance in their respective cell type were checked mainly using publicly available knowledge bases that are orthogonal to the functional annotation sources used in computing functional similarities, including ZFIN (The Zebrafish Information Network^10^), OMIM (Online Mendelian Inheritance in Man^11^), NCBI gene summaries, and PaperBLAST^12^. Notes from such databases and literature were kept for each gene to facilitate repeated inspections during manual curation.

#### Mutagenesis by CRISPR-cas9

Guide RNAs (gRNAs) were designed using the online CRISPR design tools CHOPCHOP^13,14^, as well as the web tools from IDT (Integrated DNA Technologies). Three gRNAs with high scores from both CHOPCHOP and IDT were chosen for each gene (table S5). Cas9 protein, crRNAs and tracrRNAs were ordered from IDT Alt-R CRISPR-Cas9 System. crRNAs and tracrRNAs were duplexed to form the gRNAs, which were then complexed with Cas9 protein. RNPs are formed for each gRNA separately; RNPs targeting the gene(s) of interest were then mixed equimolarly for injection. The preparation of crRNA, tracrRNA, Cas9 protein stocks, and formation of RNA duplex and gRNA+Cas9 RNA protein complex (RNP) were done following the protocols detailed in ^15^. For RNP injections, embryos from wild-type (Tupfel longfin/AB) or *creb3l1* homozygous mutant crosses were collected 10-15 minutes post fertilization and transferred into an agarose coated plates with lanes that hold the embryos. Up to 2 nl of RNP was injected into each 1-cell stage embryo through chorion.

To generate stable mutants for *creb3l1*, the founders were fin-clipped around 1 month of age, and genotyping PCR was performed using primers flanking the gRNA targeting regions. We designed primer pairs such that the product sizes are below 250 bp and the primers contain the indices compatible with illumine Mi-seq sequencing chemistry. We sent the genotyping PCR products for sequencing to select for the founders with desired frameshift mutations. The selected founders were then crossed with wild-type fish after reaching sexual maturity (> 2 month old) and the F1s were genotyped. The F1 fish that were heterozygotes with desired mutations were then in-crossed, and their offspring genotyped to select for homozygous mutants. We were not able to obtain homozygous loss-of-function mutant for *xbp1*, as they all died within 1 month post-fertilization. Similarly, we were unable to maintain a *creb3l2* homozygous mutant line.

#### Immunostaining of Hatching Enzyme and Collagen

Ctsl1b antibody were purchased from GeneTex, Inc. (North America), product catalog number: GTX128324. Collagen II antibody were purchased from abcam, product catalog number: AB34712. Both antibodies are polyclonal raised in rabbit.

Embryos were cultured to 28 hpf and dechorionated as described previously (see Methods for chapter 2). Dechorionated embryos were fixed at 4% formaldehyde at 4°C overnight and passed through a methanol dehydration series (rinsed 2x 5 min with PBST (PBS + 0.1% Tween-20), 5 min 1:1 methanol: PBST, 2 x 5 min methanol). Dehydrated embryos were kept at −20° at least overnight to permeabilize. Prior to immunostaining, embryos were rehydrated (5 min 1:1 methanol:PBST, 2 x 5 min PBST). An optional antigen retrieval step was performed, which entails washing the embryos in Tris buffer for 5 min (150 mM Tris–HCl, pH 9.0), followed by equilibrating the embryos in fresh Tris buffer at 70 °C for 10-15 min. Blocking was then performed by incubating embryos in blocking solution (10% NGS and 1% BSA in PBST) for 2-3h at RT (room temperature), or overnight at 4 °C. Primary antibodies were diluted in the blocking solution (1:100 for the collagen II antibody and 1:250 for the ctslb antibody). Blocked embryos were incubated in the diluted primary antibody solution overnight at 4 °C, on a rocking table.

The next day, embryos were washed with PBST (6 x 15min) and then incubated in the secondary antibody solution (anti-rabbit with Alexa Fluor 647, diluted 1:500 in 1% BSA in PBST) for 2-4h at RT or overnight at 4 °C in dark. Embryos were then washed extensively with PBST (6 x 15 min) before DNA and membrane staining. A staining solution was prepared with 1:1000 dilution of DAPI and WGA (Wheat Germ Agglutinin, conjugated with Alexa Fluor 488) in PBST. Embryos were incubated in the diluted staining solution for 2-4h at RT or overnight at 4 °C, washed in PBST (2x15min), and stored in PBST at 4°C before imaging.

Imaging was performed within one week after completion of immunostaining. Embryos were mounted in 2% low-melting point agarose for imaging. Imaging was performed with Zeiss LSM 880 confocal microscope using a 40x objective.

#### Hatching Rate Assay

Clutches of embryos were collected and cultured in 10cm petri-dishes up to 24hpf. Healthy looking embryos were then sorted into 24-well plates with 1 embryo per well. Each well is filled with 1.5ml of blue water (E3 medium with 0.1% methylene blue as a fungicide). Plates of embryos were kept in a 28°C incubator in dark. Hatching events were counted once each day until 6dpf without changing the medium. Time spent during counting is minimized and plates were handled gently in order to reduce mechanical perturbations and temperature changes that might affect hatching.

#### Culturing and Collecting Embryos

Embryos from wild-type (Tupfel longfin/AB) crosses were collected 20 minutes after fertilization. They were dechorionated by incubation in 1 mg/ml pronase (protease from Streptomyces griseus, Sigma-Aldrich) for 5–6 minutes until chorions began to blister, submerged in ∼200ml of zebrafish E3 embryo medium (5 mM NaCl, 0.17 mM KCl, 0.33 mM CaCl2, 0.33 mM MgSO4, dissolved in fish system water, pH to 7.3) in a glass beaker, and then the E3 medium was decanted and replaced 3 times. It was critical to prevent embryos from contacting air or plastic during this process or at any point thereafter until epiboly completes. Embryos were then cultured at 28°C in blue water in plastic Petri dishes that had previously been coated with 2% agarose (dissolved in blue water).

#### scRNA-seq Data Acquisition and Preprocessing

Embryos from wild-type (Tupfel longfin/AB) or *creb3l1* homozygous mutant crosses were collected, injected, dechorionated, and staged. Single cell suspensions were obtained using an embryo fractionation protocol: ∼30 dechorionated embryos for each experimental condition were transferred into a 2 ml LoBind Eppendorf tube (EP-022431048) filled with 1.5 ml chilled deyolk buffer (55 mM NaCl, 1.8 mM KCl, 1.25 mM NaHCO3). Embryos were pipetted up and down gently with a p1000 tip 3-4 times and placed in a Thermal shaker at 1200 rpm at 4°C for 20 sec. Tubes were then spun down at 4°C with 250g for 4min. Supernatant was discarded and 1 ml deyolking wash buffer (10 mM Tris pH 8.5, 110 mM NaCl, 3.5 mM KCl, 2.7 mM CaCl_2_) was added to each tube to resuspend the pelleted bruised embryos. Resuspension was down by pipetting 6 times with a p1000 tip. The tubes were then centrifuged again, supernatant discarded, and cell pellets resuspended with 1 ml DMEM (gibco, 11594426) with 0.1% BSA. Centrifugation and resuspension with DMEM (with 0.1% BSA) were repeated once. Tubes were then centrifuged once more. Supernatant was discarded and the final cell pallet was resuspended with 300 – 500 μl DMEM with 0.05% BSA. Cell suspension was mixed 20 times with a p200 tip and passed through a 40 μm cell strainer (Flowmi Cell Strainer, Bel-Art, H13680-0040). Cell concentration and viability in the resulting cell suspension was then checked on a hemocytometer using AO/PI viability dye (0.002% acridine orange and 0.02% propidium iodide).

Single-cell transcriptomes were generated from the single cell suspensions using 10x Chromium Single Cell 3’ V3 or V3.1 chemistry, following standard 10x protocol. For libraries prepared with 10x Chromium Single Cell 3’ V3.1 chemistry, several modifications were made to the dissociation protocol described above to improve cell viability and integrity. Briefly, after being transferred to 2ml Eppendorf tubes, embryos were first incubated in a protease solution (10 mg/ml BI protease [Sigma, P5380], 125 U/ml DNaseI, 2.5 mM EDTA in DPBS) on ice for 4min. Embryos were then washed with a stop solution (30% FBS, 0.875 mM CaCl2 in DPBS) and gently triturated (by pipetting) in Ringer’s solution. Centrifuging, resuspending, and filtering steps were then performed as in the previous protocol, but with the Ringer’s solution used as the deyolking wash buffer.

Sequencing libraries were sequenced on Illumina Nova-seq platform. Libraries were sequenced 2-3 times to achieve desired sequencing depth. The resulting reads were de-multiplexed and aligned against the Ensembl Zv11 Release 99 reference genome using *CellRanger*. The data was then filtered to exclude 1) low complexity transcriptomes with low number of genes and a total UMI counts, 2) transcriptomes with high mitochondrial gene content (>15%), and 3) transcriptomes with ectopic number of reads and are thus more likely to be doublets. A summary of the scRNA-seq samples used in our analyses can be found in Table S6.

#### scRNA-seq Batch Effect Correction and Cell Type Clustering

After obtaining cells that passed the quality control filters, we normalized the read counts in the cells such that the counts sum up to 15,000 for each cell. A value of 1 was added to each entry in the data matrix, and the resulting data was transformed to log2 space for downstream analysis. Our data for crispants were collected in 3 batches at two different institutes (batch 1 collected at the Bauer Core, Harvard University, USA, batch 2, 3, collected at Biozentrum, University of Basel, Switzerland). A small batch effect was observed between samples collected at the two different institutes. We used the external.pp.bbknn() function in the Scanpy python module with default settings to remove such batch effects^16^. The function is based on a Batch Balanced K-Nearest Neighbor (BBKNN) algorithm^17^ and projects data from different batches onto the same embedding for data visualization and clustering analysis without modifying values in the expression data matrices. UMAP and clustering were performed on the batch corrected data embedding. Clustering was done using the tl.leiden() function from the Scanpy module with parameter resolution=1.

The above analyses were performed using only the highly variable genes. Highly variable genes were found using two different methods: 1) as described before for finding variable genes in 12-24hpf scRNA-seq data, by fitting a null model (negative binomial distribution) to the raw count data to explain the expected technical variation for each gene. Genes were considered highly variable if their coefficient of variation (CV) sufficiently exceeded the null model (CV >= 0.5); and 2) using the pp.highly_variable_genes() function from the scanpy module, which identifies highly variable genes as the ones exceeds thresholds in both log mean normalized expression (>= 0.25) and normalized dispersion (variance/mean >= 0.5). Variable genes found with the two approaches were combined to be used as the highly variable genes in our analysis.

Marker genes for each cell cluster were identified as those expressed more highly in the cluster compared to the rest of the cells using the Wilcoxon test. The expression profiles of the significant marker genes with high fold changes were then checked on ZFIN and on our previously built developmental trajectories to annotate the cell types the clusters represent.

#### Endogenous Target Gene Inference by Linear Regression

Target inference were done by performing linear regression to model each gene detected across single cells in a cell type as a linear combination of a constant value (baseline expression) and the UPR TF genes in the same cells. Prior to regression, UPR TF levels used as the predictor variables were adjusted, such that a TF’s level is set to 0 in the samples where it was mutated or targeted by CRISPR. UPR TFs themselves were excluded from the response variables – the genes whose expressions were to be predicted using the UPR TFs. Two additional predictors were added to the predictor variables: 1) “injection”, which was set to 0 for cells from uninjected TLAB and *creb3l1*-mut (uninjected *creb3l1* homozygous mutant) embryos, set to 1 for cells from embryos injected with gRNAs targeting 1 gene, and set to 1.5 for cells from embryos injected with gRNAs targeting multiple genes; 2) “batch”, which was set to 0 for the transcriptomes collected at Harvard, and set to 1 for those collected at Biozentrum. Injection was included as a predictor variable because we previously observed that DNA breaks induced by Cas9 could induce genes involved in apoptosis in some cells. Injection and batch were added as predictors so that expression changes due to injection or batch are less likely to be confounded with the changes caused by loss of UPR TF functions.

Regression was then performed using a robust linear model implemented by the RLM() function provided by the python module statsmodels (v0.11.0)^18–21^. AndrewWave function was specified as the robust criterion function for downweighing outliers by setting the parameter “M = sm.robust.norms.AndrewWave()”. For each response variable (potential target genes), the RLM function returns a linear model that estimates the level of the response variable as a linear combination of the predictor variables (TFs, injection, batch). A coefficient and a p-value is calculated for each predictor variable. An R^2^ score (coefficient of determination) was calculated for each gene to inform how well the variance in the gene is predicted by the model. The R^2^ score is calculated as 𝑅^2^ = 1 −^𝑆𝑆^𝑟𝑒𝑠⁄^𝑆𝑆^*tot*, where 𝑆𝑆_𝑟𝑒𝑠_ denotes the sum of squared residual values (i.e. predicted – real gene expression value), and 𝑆𝑆_𝑡𝑜𝑡_ denotes the sum of squared differences from the mean (i.e. expression in a cell – mean expression in the cell type). The coefficients, p-values, and R^2^ scores were thresholded to call target genes (R^2^ > 0.15, coefficient > 0.05, p-value < 1e-9). Thresholds were picked by varying the values while checking the expression profile of included and excluded target genes across samples. A relatively stringent threshold was picked such that the vast majority of the inferred targets are visually differentially expressed in the corresponding samples compared to the control on a heatmap.

#### In Vitro Transcription of UPR TF mRNAs

Sequences of the UPR TF cDNAs were obtained from NCBI. Primers were designed to amplify the desired cDNA sequences. For *xbp1*, primers flanking the entire coding sequence for *xbp1s* were used. For *creb3l1*, *creb3l2* and *atf6*, the protein sequences were first aligned to their mammalian orthologs to identify the transmembrane domains, then primers were designed to amplify the cDNA sequence coding only the cytoplasmic domains. For *creb3l2*, an additional primer pair was designed to amplify the whole coding sequence. The primers were designed with 30 nt that matches the pCS2 plasmid sequence flanking the insertion site, so that the PCR products can be combined with the plasmid backbone by Gibson assembly (table S4.2). cDNAs generated from zebrafish embryos at 4 hpf, 5.3 hpf, 6 hpf, 8 hpf, 12 hpf, 24 hpf and 48 hpf were pooled together for amplification of the UPR TF cDNA sequences. Gibson assembly were used to integrate PCR amplified cDNA sequences with the PCR amplified pCS2 backbone. α-select gold competent cells were transformed with the integrated plasmids and grew at 37 °C to form individual colonies. Individual colonies were further grown, from which plasmids were isolated through miniprep (Plasmid DNA mini kit, EZNA) and sent for Sanger sequencing. The plasmids harboring the desired insertions (UPR TF coding sequences) were linearized by incubation with restriction enzymes (NEB). Linearized plasmids were subjected to invitro transcription using a SP6 Transcription Kit (Invitrogen AM1340). The resulting mRNA was purified (Total RNA Kit I, EZNA), quantified (nanodrop), and 2 μl was denatured (diluted in 8 μl nuclease free water and incubated at 70 °C 10min) then checked on an agarose gel. mRNAs with concentrations greater than 200 ng/μl and appeared as one single band on the agarose gel were deemed with good quality. mRNAs with good quality were stored as 200 ng/μl aliquots at -80 °C. mRNAs were thawed and diluted to desired concentrations 1-0.5 hours before performing injection for the overexpression experiments.

#### Mis-expression of UPR TFs

Embryos from wild-type (Tupfel longfin/AB) crosses were collected 10-30 minutes post fertilization. For *mCherry* and *creb3l2*-full overexpression, embryos were injected with 125pg and 40 pg of the corresponding mRNAs, respectively. For *atf6N*, *creb3l1N*, and *creb3l2N* mis-expressions, embryos were first injected with Cas9 protein complexed with *xbp1* targeting gRNAs. 40pg mRNAs encoding *atf6N*, *creb3l1N*, *creb3l2N*, or *mCherry* were then injected before cells entered the 2-cell stage. For *xbp1*-s and *bhlha15* mis-expressions, *xbp1*+*atf6* double crispants were first generated by injecting Cas9 protein and gRNAs. 70pg *xbp1*-s mRNA and/or 50pg *bhlha15* mRNA, or 120pg *mCherry* mRNA were then injected before cells entered the 2-cell stage. All embryos were injected through chorions. The total volume injected in each embryo was controlled below 2.5nl.

#### Cell Type Identification for Mis-expression Data

Mis-expression of the UPR TFs induced global transcriptional shifts between samples (Figure 5B, Figure S7B). To identify consistent biological cell types, we first analyzed the *mCherry* control dataset and defined variable genes, performed PCA, UMAP, and Leiden clustering as described previously for the crispant dataset. Each of the other datasets were projected onto the *mCherry* data embedding using the tl.ingest() function from scanpy, such that cells’ UMAP coordinates and cell cluster memberships are mapped according to their similarity to the control cells in the principle component space^16^. The expression data matrices remain unchanged after the ingest operation.

Marker genes for each cell cluster were identified as those expressed more highly in the cluster compared to the rest of the cells using the Wilcoxon test. The expression profiles of the significant marker genes with high fold changes were then checked on ZFIN and on our previously built developmental trajectories to annotate the cell type each cluster represents.

#### Induced Target Gene Inference by Linear Regression

Induced target genes were inferred using the mis-expression data. Target inference was performed using the same method as for endogenous target gene inference with a few modifications: 1) UPR TF levels in the sc-transcriptomes was used as the predictor variables since injected mRNAs were quantitatively detected in scRNA-seq. 2) A regression model was built for each UPR TF separately using the samples mis-expressing that UPR TF and the appropriate control sample. For instance, models that predicts *xbp1-s* regulation were built using *xbp1-s* and *mCherry* mis-expression samples, with each gene’s expression levels modeled as g=bias+C\**xbp1-s*, where g is the expression of the gene, bias reflects baseline expression level, and C represent the regulatory activity (coefficient) of *xbp1* on the gene.

## Supplemental Tables

**Table S1.** Significantly enriched GO terms among notochord-enriched genes. This table is provided as an Excel file.

**Table S2.** Significantly enriched GO terms among hatching gland-enriched genes. This table is provided as an Excel file.

**Table S3.** List of gene modules in the notochord. This table is provided as an Excel file. Genes with mentions in the main text are underscored. Modules are ordered alphabetically. Genes without module assignment are in the “Unassigned” group as the last row of the table.

**Table S4.** List of gene modules in the hatching gland. This table is provided as an Excel file. Genes with mentions in the main text are underscored. Modules are ordered alphabetically. Genes without module assignment are in the “Unassigned” group as the last row of the table.

**Table S5.**
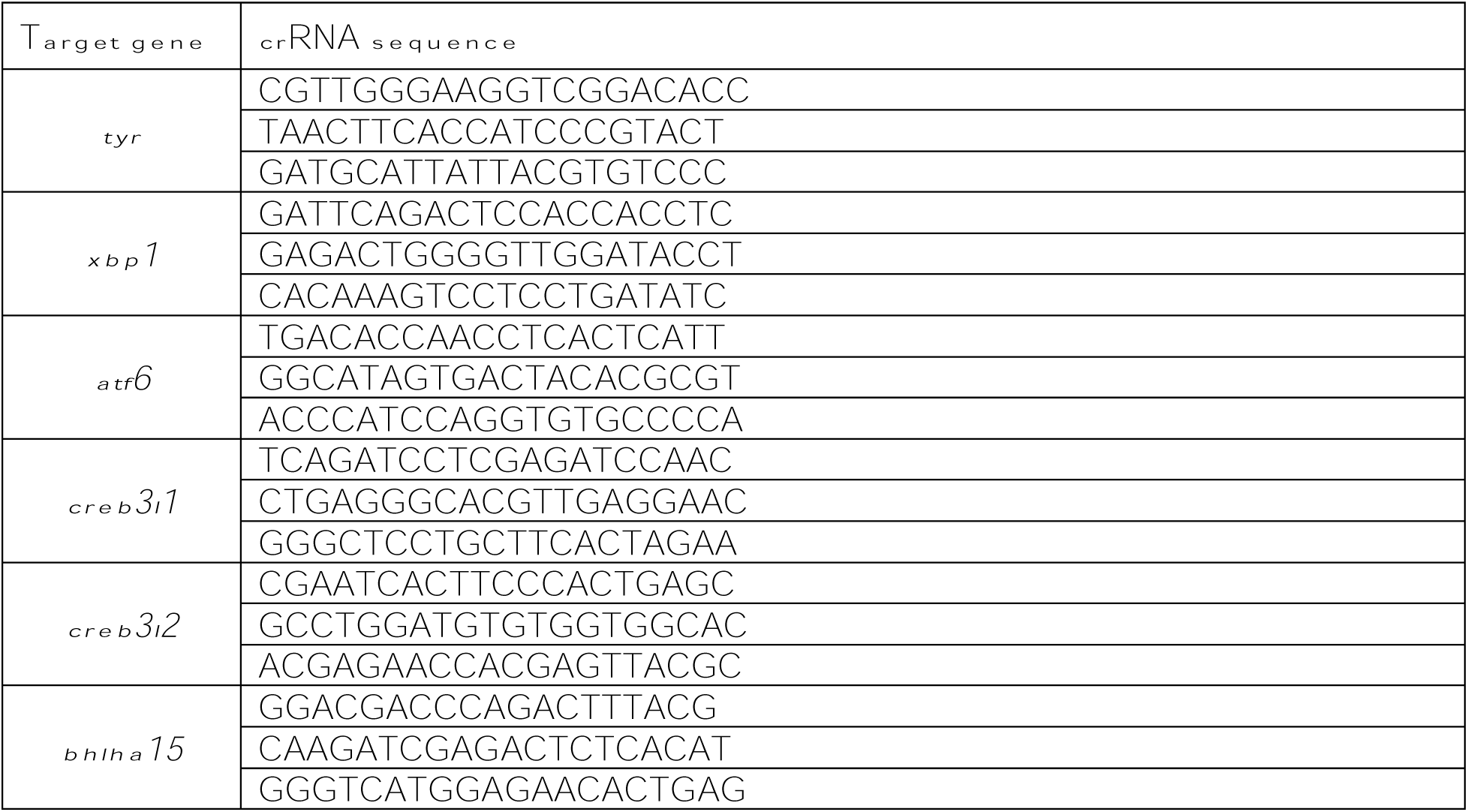
Guide RNA sequences.

**Table S6.** Summary of the scRNA-seq datasets used in UPR TF target gene inference. This table is provided as an Excel file.

**Table S7.** Lists of UPR TF target genes. This table is provided as an Excel file. Only positive targets are included. Endogenous targets were inferred from the loss-of-function dataset in each cell type. Induced targets were inferred from the mis-expression dataset; genes induced in at least 2 cell types were included. The general secretion, ECM secretion, and gland-like secretion programs were defined in the main text (see section “*creb3l1*/*3l2* and *xbp1* regulate ECM secretion and gland-like secretion programs, respectively”).

## Notes

### Competing Interest Statement

The authors have declared no competing interest.

