## Supplementary material for "Gene module reconstruction elucidates cellular differentiation processes and the regulatory logic of specialized secretion": Figure S

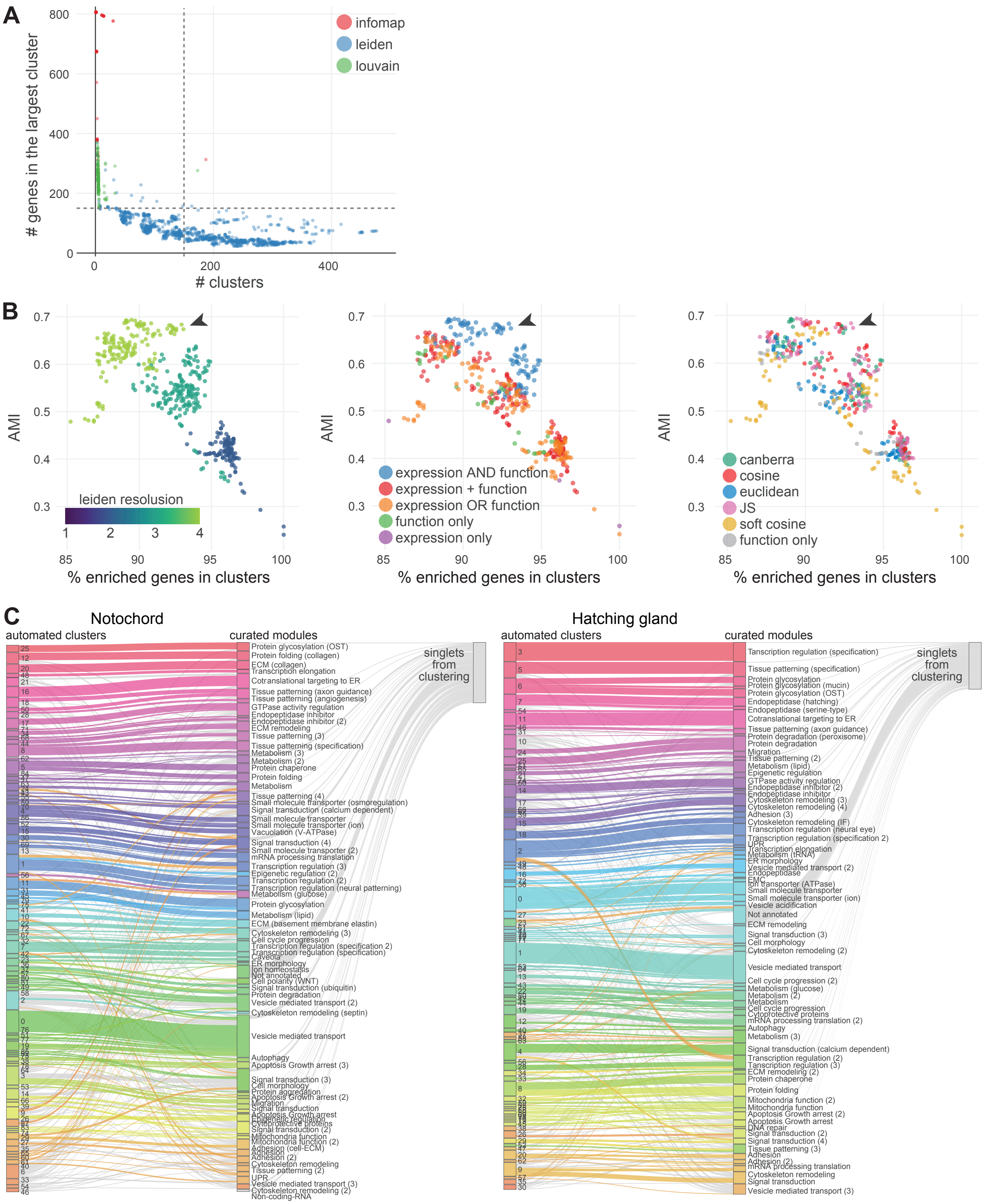


**Figure S1. Comparison between automated clusters and manually curated modules.**

**Figure S1. Comparison between automated clusters and manually curated modules. A)** Plot of the number of clusters and the size of the largest cluster resulted from the various clustering methods tested for the notochord-enriched genes. Each point represents the clustering result from a clustering method (clustering algorithm and similarity metric) colored by the algorithm used. Methods that produced <150 clusters (left to the vertical dashed line) with the largest cluster containing <150 genes (below the horizontal dashed line) were selected for further evaluation. **B)** Percent of notochord-enriched genes included in non-singlet clusters (x-axis) and the agreement between manually curated modules (from the subset of 300 enriched genes) and automated clusters (measured by AMI, y-axis) were used to further select the best clustering method. Only methods passed the criteria in A) were considered. Arrowhead marks the method chosen to produce the final clusters. **C)** Comparison between clusters produced with the method selected in B) and the final gene modules after manual curation. Ribbons represent member genes.

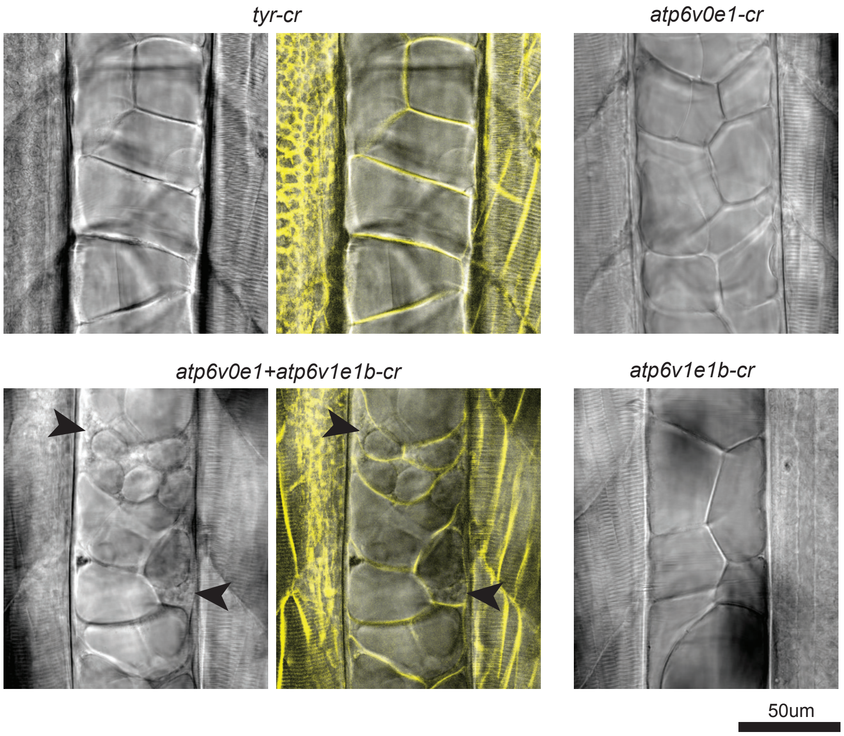


**Figure S2. *atp6v0e1* is involved in maintaining notochord vacuole structures.** Crispants were generated by injecting gRNAs into 1 cell stage transgenic embryos with mCherry-CAAX. Crispant notochords were imaged in 3dpf (days-post-fertilization) live embryos. Both DIC (grey scale) and mCherry-CAAX fluorescence (yellow, cell membrane) were recorded. The control *tyr* crispant (cr) notochord contains big vacuoles that filled up entire notochord cells, while *atp6v0e1* + *atp6v1e1b* double crispants exhibited fragmented vacuoles (arrow heads). Single crispants of *atp6v0e1* or *atp6v1e1b* did not show obvious abnormality in notochord vacuoles.

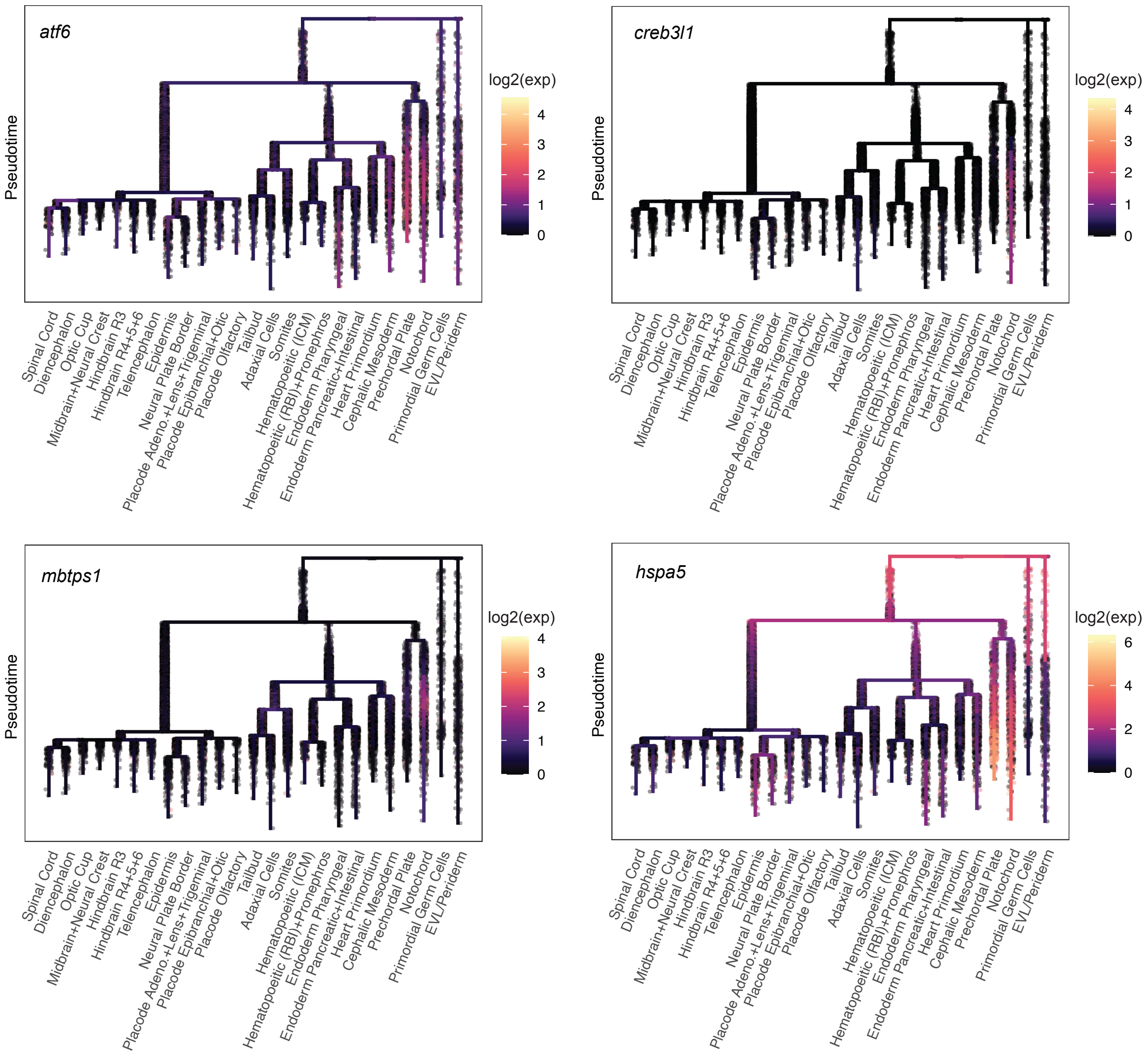


**Figure S3. UPR transducer genes are specifically enriched in the axial mesoderm.** In each plot, expression level of a UPR transducer gene is overlaid on the zebrafish developmental trajectory spanning 3-12hpf. Development progresses from top to bottom. Expression of *xbp1* and *creb3l2* are included in **Figure 3A**.

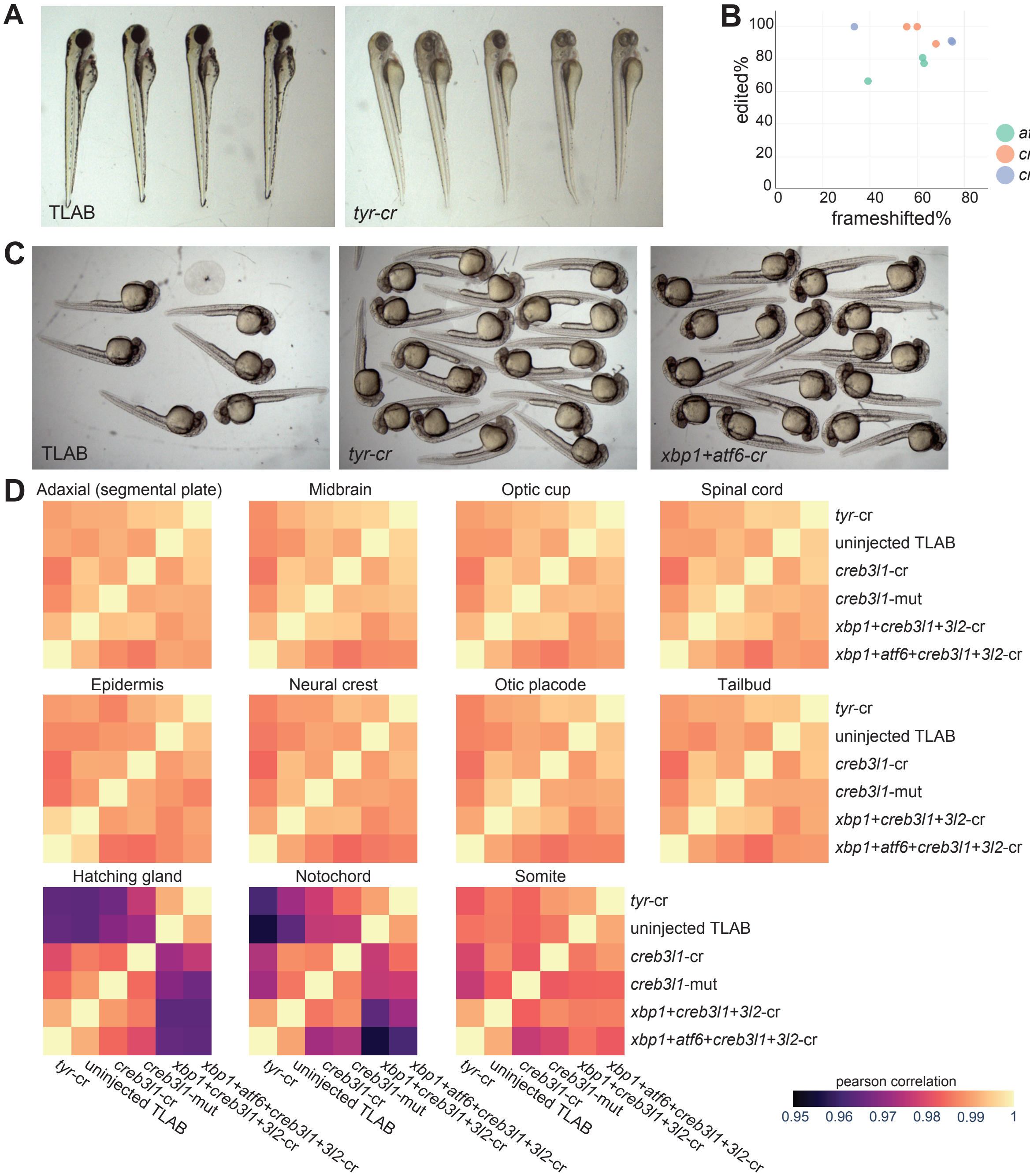


**Figure S4. CRISPR/Cas9 is highly effective in generating specific F0 loss-of-function phenotypes. A)** Representative uninjected TLAB (wild-type) and *tyr* crispants at 3dpf. 30 out of 30 *tyr* crispants exhibited loss of or greatly reduced pigmentation. **B)** Mutagenesis rate of individual gRNAs used for generating *atf6*, *creb3l1*, and *creb3l2* crispants. 3 gRNAs for each gene were assessed individually. Genotyping PCR products were sequenced with MiSeq and analyzed for mutation rates using the R package ampliCan. All 3 gRNAs for each gene were coinjected to generate the crispants used in the study. **C)** Uninjected TLAB, *tyr* crispants, and *xbp1+atf6* crispants at 28hpf. Crispants showed no gross developmental defects. **D)** Heatmaps showing pearson correlations among different scRNA-seq samples for several cell types. The 12hpf scRNA-seq data from one experimental batch were used. Correlations were calculated on the log mean expression of the 1594 highly variable genes per sample per cell type. Samples in all heatmaps follow the same order. In most cell types, crispant and non-crispant samples highly correlated with each other, while in the notochord and hatching gland, *xbp1+creb3l2+creb3l1*-cr and *xbp1+atf6+creb3l2+creb3l1*-cr showed lower correlation with other samples, consistent with the tissue-specific expression and phenotypes observed. *xbp1+creb3l2+creb3l1*-cr and *xbp1+atf6+creb3l2+creb3l1-cr* embryos were generated by injecting Cas9 and gRNAs targeting *xbp1* and *creb3l2* (and *atf6*) in the *creb3l1* homozygous mutant embryos. “*creb3l1*-mut”: *creb3l1* homozygous mutant.

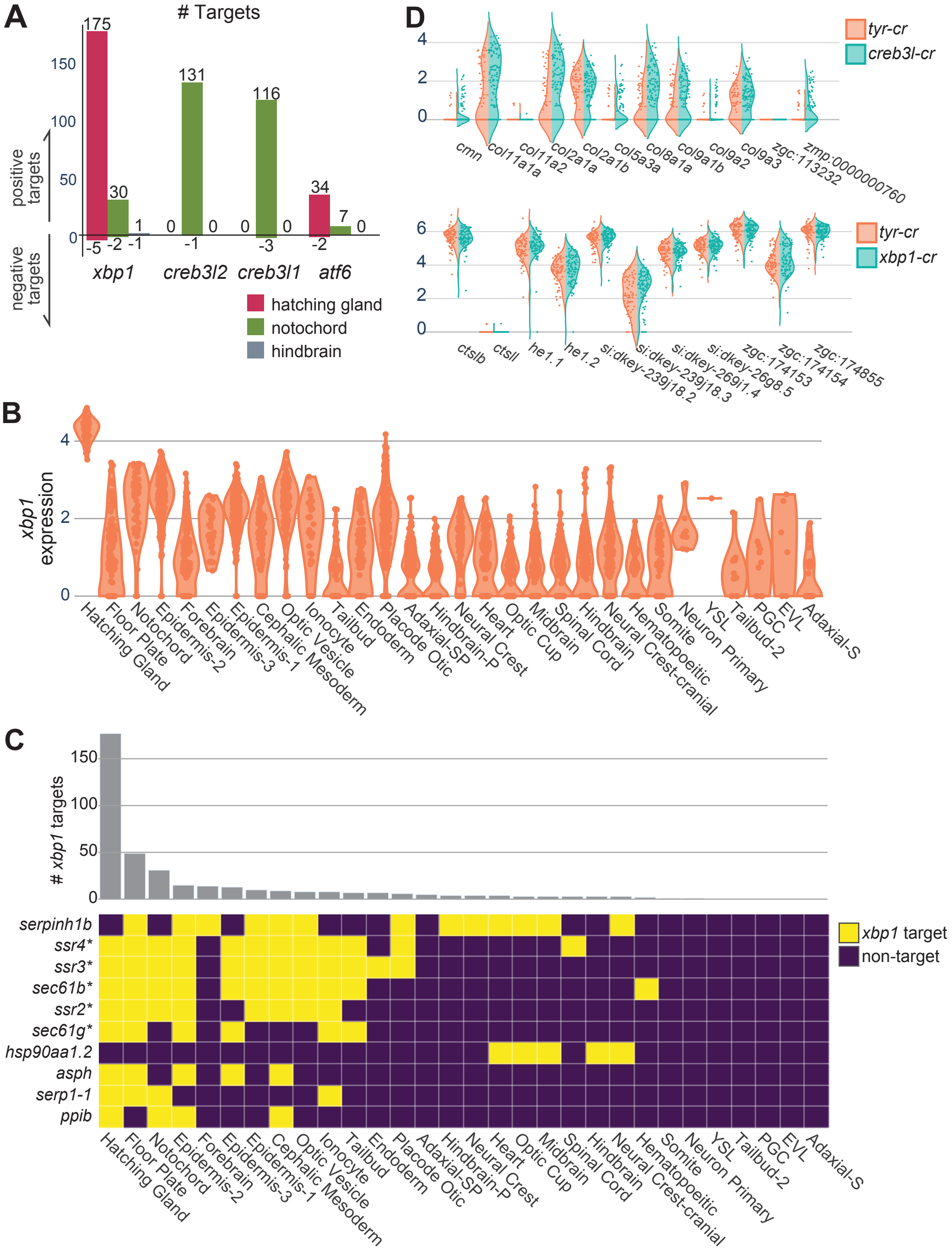


**Figure S5. Endogenous UPR TFs are active primarily in the axial mesoderm as transcriptional activators. A)** Number of endogenous target genes identified from the loss-of-function analysis for each UPR TF in the hatching gland, notochord, and hindbrain. Hindbrain serves as a representative non-secretory cell type. **B)** Log expression level of *xbp1* in each cell type in the control crispant (*tyr*-cr) at 12hpf. **C)** *xbp1* regulates a small set of sensitive targets in several cell types. Upper: number of *xbp1* positive targets across cell types. Lower: genes identified as a target in more than 3 cell types. Genes marked with “*” are involved in protein co-translational targeting to the ER. These genes are regulated by *xbp1* in several cell types in addition to the hatching gland, suggesting that they respond to even low levels of *xbp1* activity. **D)** Secretory cargos are not dysregulated in UPR TF crispants. Upper: expression of notochord cargos (collagens) in the notochord. Left violin: *tyr* crispant; right violin: *creb3l1+3l2* crispant. Lower: expression of hatching gland cargos (endopeptidases for hatching) in the hatching gland. Left violin: *tyr* crispant; right violin: *xbp1* crispant.

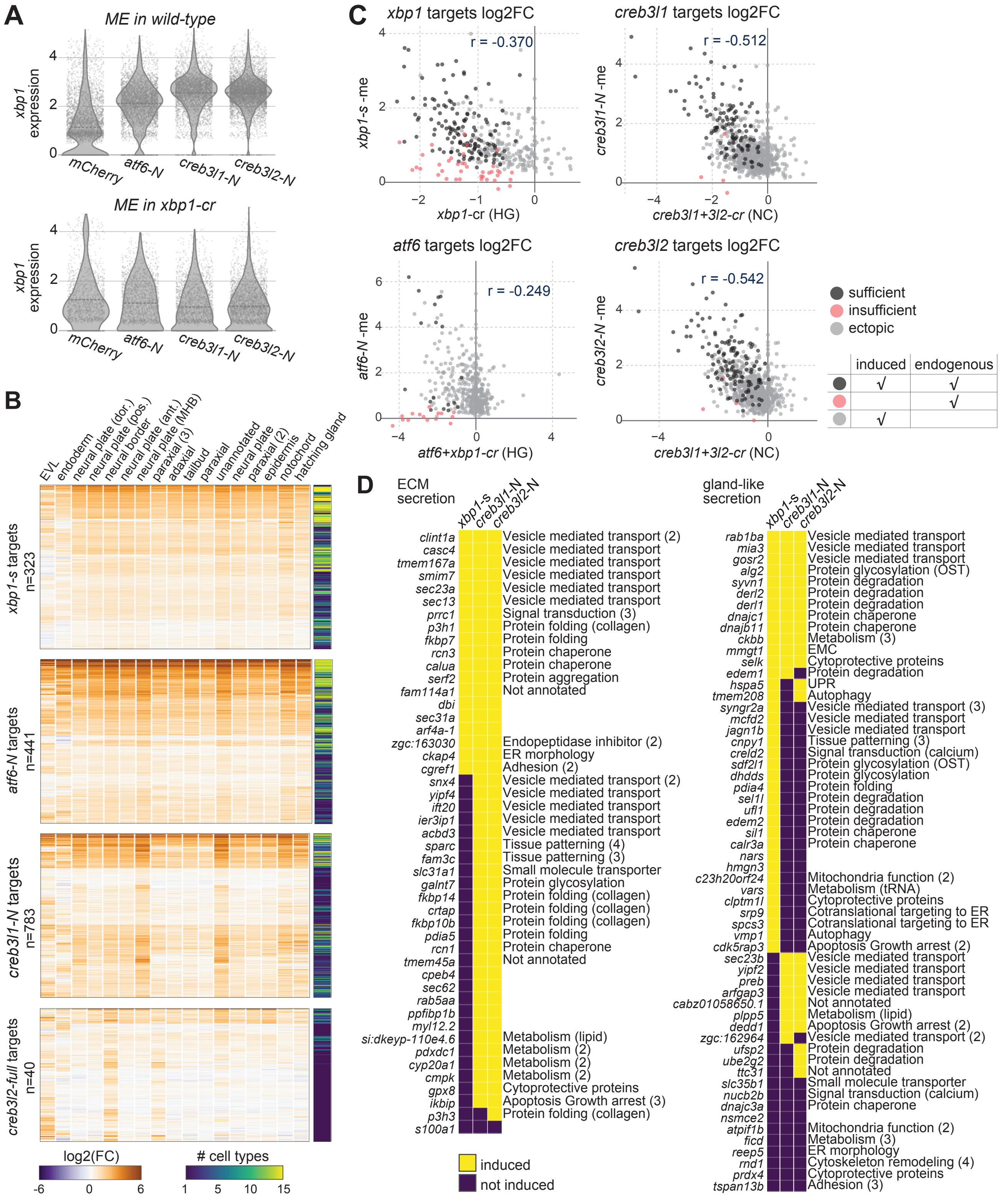


**Figure S6. Mis-expressed UPR TFs induced hundreds of genes in the early embryo.**

**Figure S6. Mis-expressed UPR TFs induced hundreds of genes in the early embryo. A)** *xbp1* is a target of the other UPR TFs. *xbp1* expression (log scaled) are shown as violin plots. Each point represents a single-cell transcriptome. Upper: *xbp1* expression is induced by *atf6-N*, *creb3l1-N*, or *creb3l2-N* mis-expressions. Lower: induction of *xbp1* was mitigated when UPR TFs were mis-expressed in *xbp1* crispants. **B)** Log2 fold changes of UPR TF mis-expression induced target genes (rows) by cell types (columns). Bars on the right show the number of cell types in which a gene is identified as a target. Result for *creb3l2*-N targets is shown in Figure 5C. **C)** Target gene fold changes in mis-expression and crispant samples. Y-axes: mean log2 fold change in mis-expression samples compared to *mCherry* control across cell types. X-axes: log2 fold change in crispants compared to *tyr-cr* control in indicated cell types (HG: hatching gland; NC: notochord). Each point is a target gene colored according to their regulation. The union of mis-expression induced and endogenous targets are plotted for each UPR TF. Only positive targets are shown. Pearson correlations between the log2 fold changes in mis-expressed and crispant samples are labeled on the plots. The correlation is stronger for *creb3l1* and *creb3l2* targets and is the weakest for *atf6 targets*. **D)** Identities of ECM secretion (left) and gland-like secretion (right) genes and their induction by *xbp1-s*, *creb3l1-N*, and *creb3l2-N*. Gene symbols are on the left of the heatmaps and module memberships on the right. Genes without module membership are not identified as enriched genes from the trajectory data.

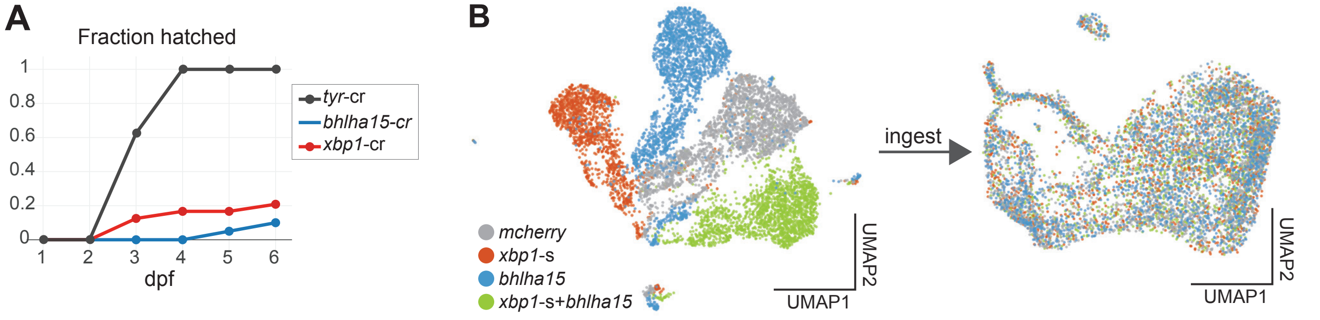


**Figure S7. *bhlha15* loss- and gain-of-function phenotypes. A)** *xbp1* and *bhlha15* crispants exhibited reduced hatching. Number of embryos assayed for each condition: *tyr*-cr = 8; *xbp1*-cr = 24; *bhlha15*-cr = 20. **B)** UMAP of *xbp1-s* and *bhlha15* mis-expression scRNA-seq data colored by mis-expression conditions. Left: UMAP projection before dataset integration. Right: UMAP projection of the same data integrated with the ingest algorithm with the control data (*mCherry*) as the reference.
